# Coordinated DNA and Histone Dynamics Drive Accurate Histone H2A.Z Exchange

**DOI:** 10.1101/2021.10.22.465479

**Authors:** Matthew F. Poyton, Xinyu A. Feng, Anand Ranjan, Qin Lei, Feng Wang, Jasmin S. Zarb, Robert K. Louder, Giho Park, Myung Hyun Jo, Joseph Ye, Sheng Liu, Taekjip Ha, Carl Wu

**Affiliations:** Department of Biophysics and Biophysical Chemistry, Johns Hopkins School of Medicine, Baltimore, Maryland, USA; Department of Biology, Johns Hopkins University, Baltimore, Maryland, USA; Department of Biophysics, Johns Hopkins University, Baltimore, Maryland, USA; Laboratory of Biochemistry and Molecular Biology, Center for Cancer Research, National Cancer Institute, Bethesda, Maryland, USA; Department of Biomedical Engineering, Johns Hopkins School of Medicine, Baltimore, Maryland, USA; Howard Hughes Medical Institute, Baltimore, Maryland, USA; Department of Molecular Biology and Genetics, Johns Hopkins School of Medicine, Baltimore, Maryland, USA

## Abstract

Nucleosomal histone H2A is exchanged for its variant H2A.Z by the SWR1 chromatin remodeler, but the mechanism and timing of histone exchange remain unclear. Here, we quantify DNA and histone dynamics during histone exchange in real-time using a three-color single-molecule FRET assay. We show that SWR1 operates with timed precision to unwrap DNA with large displacement from one face of the nucleosome, remove H2A-H2B from the same face, and rewrap DNA, all within 2.3 seconds. Such productive DNA unwrapping requires full SWR1 activation and differs from unproductive, smaller-scale DNA unwrapping caused by SWR1 binding alone. On an asymmetrically positioned nucleosome, SWR1 intrinsically senses long-linker DNA to preferentially exchange H2A.Z on the distal face as observed in vivo. The displaced H2A-H2B dimer remains briefly associated with the SWR1-nucleosome complex and is dissociated by histone chaperones. These findings reveal how SWR1 coordinates DNA unwrapping with histone dynamics to rapidly and accurately place H2A.Z at physiological sites on chromatin.

**One-Sentence Summary:** Multicolor single-molecule FRET reveals how SWR1 unwraps DNA to exchange nucleosomal H2A-H2B for H2A.Z-H2B.

## Main Text

Eukaryotic genomes are packaged into nucleosomes which are enzymatically remodeled to change their position onDNA as well as their composition (*1*). A canonical nucleosome contains two copies of the four histones H2A, H2B, H3 and H4 (*2*). Nucleosomes can be remodeled to contain non-canonical histone variants via a chromatin remodeling event named histone exchange (*1, 3*). Histone exchange is exemplified by the removal of H2A-H2B (AB) within a canonical nucleosome (histone eviction) and its coupled replacement by the variant histone dimer H2A.Z-H2B (ZB) (histone insertion) by the one-megadalton multiprotein complex SWR1 in *S. cerevisiae* (*4–6*). SWR1 and its homologues in metazoans deposit ZB dimers into the “+1” nucleosome located immediately downstream of the transcription start site (TSS), at sites of DNA damage, replication origins and enhancer and insulator elements (*7–10*). H2A.Z regulates the efficiency of transcription by altering the stability of the +1 nucleosome which occludes the TSS (*11–14*).

Unlike other chromatin remodelers which slide nucleosomes by translocating on nucleosomal DNA while leaving the histone octamer intact, SWR1 is thought to manipulate nucleosomal DNA to destabilize the histone octamer, leading to the eviction of an AB dimer. A structural and biophysical study indicated that nucleosomal DNA is unwrapped upon SWR1 binding in the presence of ZB dimer and a non-hydrolyzable ATP analogue (*15*). SWR1 binding to nucleosomes has been shown to induce DNA unwrapping and rewrapping over a wide range of timescales from microseconds to seconds (*15, 16*), but the necessity of these unwrapping events for the eviction of an AB dimer and subsequent insertion of a ZB dimer is unknown. To directly address how DNA and histones are dynamically reconfigured during histone exchange we developed a three-color smFRET assay (single-molecule Fluorescence Resonance Energy Transfer) (*17–19*) to observe the rearrangement of nucleosomal DNA and histones in real-time.

To monitor the conformation of nucleosomal DNA and histones during histone exchange we reconstituted a nucleosome substrate with two fluorophores on DNA and one on histone H2A. A derivative of the 601-nucleosome positioning sequence (601-c2 (*20*), sequence in Table S1) was labeled with Cy5 as it ‘enters’ the nucleosome and Cy7 near the nucleosome dyad, allowing DNA unwrapping and rewrapping to be monitored via Cy5-Cy7 FRET (Fig. 1A). The C-terminal tail of H2A(K120C) was labeled with Cy3 (Fig. 1A), allowing histone exchange to be detected by the loss of Cy3 fluorescence when the Cy3-AB dimer is replaced by an unlabeled ZB dimer (Fig. 1A). The reconstituted three-color nucleosome was symmetrically positioned between two 43 bp DNA linkers (43-N-43). SWR1 remodeled the three-color nucleosome at a rate comparable to unlabeled nucleosomes in bulk (*21*) (fig. S1A) and all donor-acceptor pairs engage in FRET efficiently as designed (fig. S1C-H). Single nucleosomes immobilized on passivated quartz slides were viable substrates for histone exchange as shown by the eviction of Cy3-AB over time (Fig. 1B). All single-molecule and bulk exchange reactions were performed at room temperature, 22 °C. Exchange only occurred in the presence of ATP (Fig. 1D) and the exchange rate was sensitive to both SWR1 and ZB dimer concentrations (fig. S1I,J). In a separate experiment we were also able to visualize the insertion of Cy3-ZB into immobilized nucleosomes and validate that AB eviction and ZB insertion occur at the same rate and are tightly coupled in immobilized nucleosomes as they are in bulk (Fig. 1C,D).

**Fig. 1.**
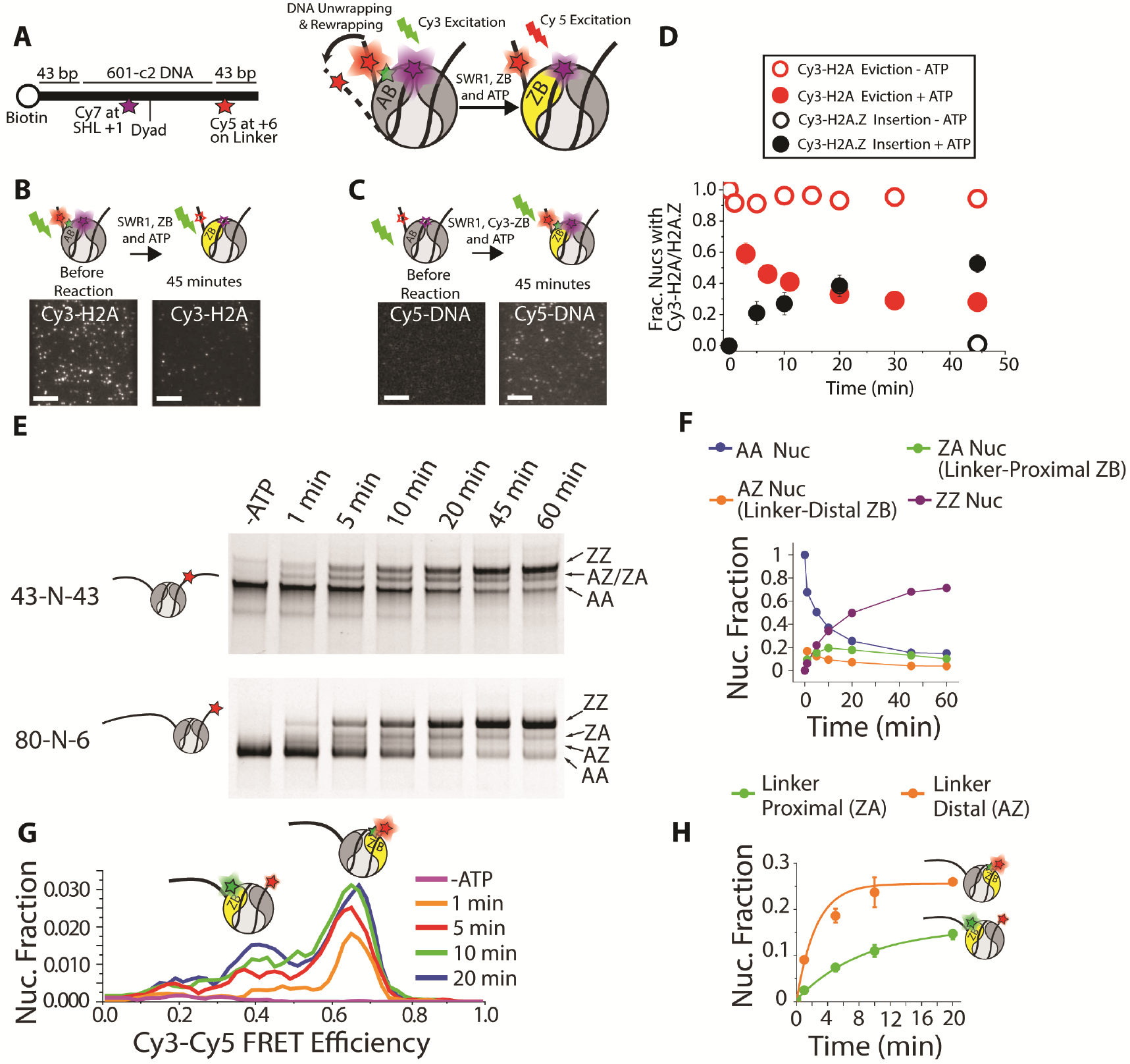
A three-color smFRET assay for histone exchange. (**A**) The nucleosome construct used to observe DNA dynamics during histone exchange. (**B**) Histone exchange detected through the loss of Cy3-Cy5 FRET spots as Cy3-AB is removed from the nucleosome. (**C**) Histone exchange detected via the appearance of Cy3-Cy5 FRET spots as Cy3-ZB is inserted into the nucleosome. Error bars are smaller than some data points. (**D**) Single-molecule histone eviction and insertion require ATP and occur at the same rate (average of three separate experiments, error is standard deviation and is smaller than datapoints for some conditions). (**E**) The EMSA assay for histone exchange utilizing 43-N-43 and 80-N-6 nucleosomes. Insertion of one or two triple-flag tagged ZB dimers causes nucleosomes to migrate slower through the gel. While the rate of histone eviction and insertion for symmetrically and asymmetrically positioned nucleosomes is the same, the linker-distal AB dimer (AZ band on bottom gel) is exchanged at a faster rate than the linker-proximal dimer (ZA band on bottom gel). The faint band below the AA nucleosome in the top gel for 43-N-43 nucleosomes likely represents a negligible amount (<5%) of hexamer contamination. (**F**) quantified band intensities from the 80-N-6 exchange reaction (as show in C). Data points are averages of two technical replicates; error bars (standard deviation) are smaller than the data points. (**G**) Imaging single nucleosomes only labeled on DNA after exchanging with Cy3-ZB in bulk confirms that the ZB is exchanged with the linker-distal dimer (high FRET population) more quickly than the linker-proximal dimer (low FRET population). (**H**) Quantified results from three replicates of single-molecule pulldown (as shown in G), showing that the linker distal dimer is exchanged four times faster than the linker proximal dimer, as determined by single exponential fits (solid lines). Error bars are standard deviation of the average.

The specific AB dimer on one face of the nucleosome exchanged for ZB can be distinguished by the appearance of high and low Cy3-Cy5 FRET populations when Cy3-ZB is inserted into a nucleosome where Cy5 is on DNA (fig. S2A-G). The single-molecule Cy3-ZB insertion time-course on the 43-N-43 nucleosome labeled only with Cy5 and Cy7 on DNA show that both high (Cy5-proximal ZB) and low (Cy5-distal ZB) FRET populations increase at the same rate (fig. S2F), demonstrating that histone exchange occurs without bias on either face of a nucleosome symmetrically positioned between equal lengths of linker DNA. However, in *S. cerevisiae,* H2A.Z is enriched on the NDR-distal (nucleosome depleted region) face of the +1 nucleosome (*22*), which is asymmetrically positioned between the NDR (∼140 bp) and the linker DNA leading to the +2 nucleosome (∼18 bp) (*23*).

To determine if the linker asymmetry could bias histone exchange, we reconstituted an asymmetric nucleosome flanked by 80 bp and 6 bp DNA linkers (80-N-6) as a mimic for the +1 nucleosome (sequence in Table S2). Given that the insertion of a Flag-tagged ZB dimer (ZB-Flag) into the nucleosome face proximal or distal to the long linker can be distinguished by electrophoretic mobility shift (*24*), we found that the ZB-Flag is inserted more rapidly into the linker-distal than linker-proximal site on 80-N-6 nucleosomes (Fig. 1E,F). Furthermore, analysis of Cy3-Cy5 FRET in single Cy5-labeled nucleosomes pulled down and imaged after exchange with Cy3-ZB dimer in bulk revealed that SWR1 exchanges ZB dimer into the long linker-distal nucleosome face at a 4-fold higher rate than the proximal face (Fig. 1G,H, fig. S3A-C)). It is well known that the 601 nucleosome positioning sequence itself can bias remodeler activity, (*24–26*). To control for sequence effects the DNA linkers were flipped to the opposite side of the nucleosome positioning sequence (6-N-80, sequence in Table S3). Quantifying the Cy3-Cy5 FRET efficiency for 6-N-80 nucleosomes after exchange at room temperature showed that Cy3-ZB was inserted into the long linker-distal face of 6-N-80 nucleosomes at a faster rate than the linker-proximal face (fig. S3D), similar to the 80-N-6 construct. These results show that asymmetric DNA linker lengths bias the histone exchange reaction to the linker-distal side of nucleosomes at near-physiological temperature. Although DNA composition, conformational flexibility, and temperature may also favor asymmetric SWR1 activity(*24, 27*), our findings indicate that sensing of long linker DNA lengths, a defining characteristic common to the vast majority of +1 nucleosomes, is intrinsically important for the asymmetric enrichment of H2A.Z on the NDR-distal face of the +1 nucleosome *in vivo*(*22*).

To examine the exchange reaction in real-time, we concurrently monitored FRET efficiencies between the three donor-acceptor FRET pairs immediately after introduction of SWR1, ATP and ZB dimer to immobilized three-color nucleosomes (imaging scheme in fig. S1B). We observed a striking increase in Cy3-H2A signal intensity lasting tens of seconds in a substantial fraction of nucleosomes (127/553) (Fig. 2A, additional examples in fig. S4A-D). In contrast, Cy3-H2A signal increase is rarely observed for nucleosomes alone (0/255), without ATP (1/47), without ZB (0/46), with SWR1 only (0/71), or with substitution of a non-hydrolyzable ATP analog (1/76) (Fig. 2B). Direct excitation shows that both Cy5 and Cy7 fluoresce continuously, indicating that the Cy3 signal increase is caused by a drop in FRET efficiency with Cy5 or Cy7, consistent with the Cy3-AB dimer being displaced from the histone octamer by SWR1 during a histone exchange reaction.

**Fig. 2.**
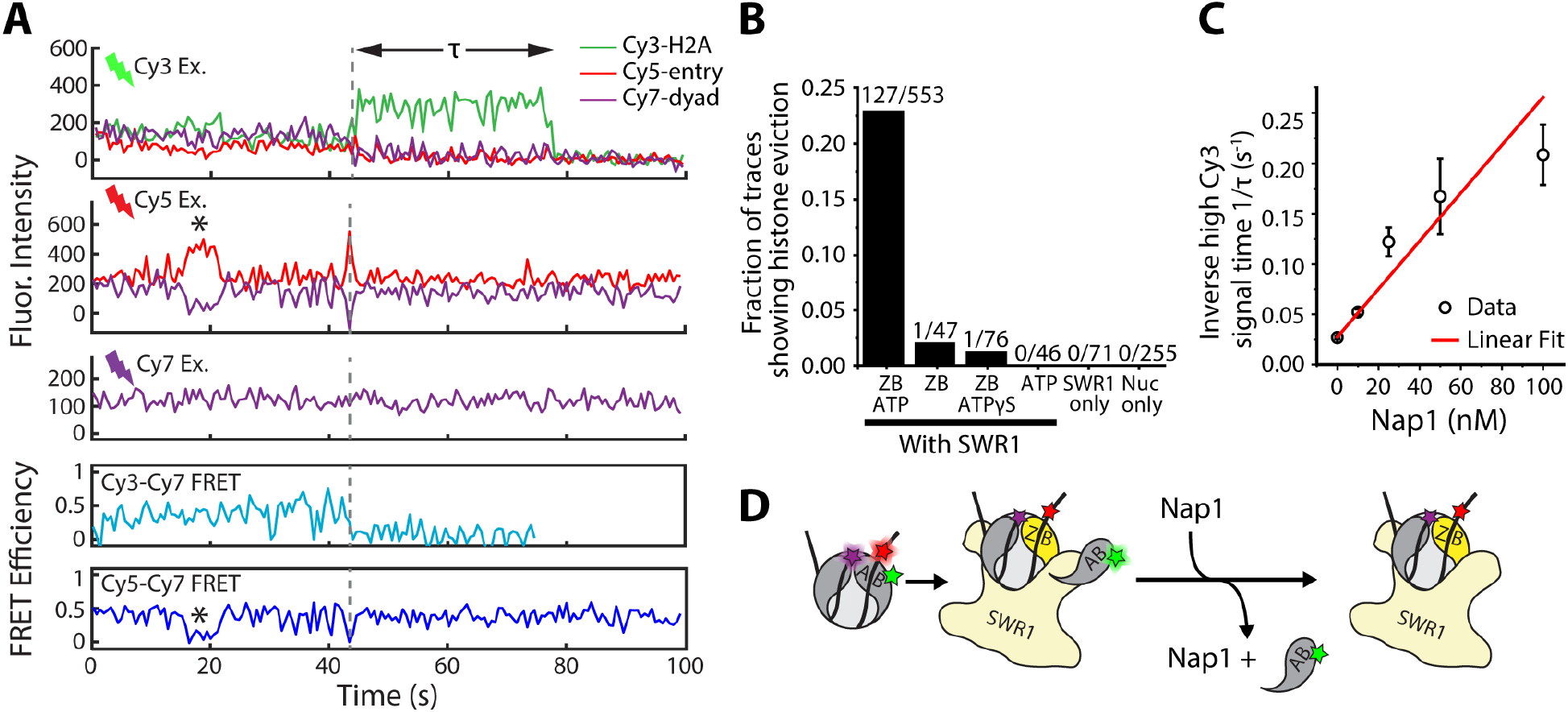
Observing histone eviction in real-time. (**A**) Representative fluorescence intensity time traces capturing histone exchange on a single immobilized nucleosome. The top three panels show Cy3 (green), Cy5 (red) and Cy7 (purple) intensities during the direct excitation of each fluorophore. The bottom two panels show time traces of the Cy3-Cy7 and Cy5-Cy7 FRET efficiencies. A grey dashed line indicates the spontaneous increase in Cy3 intensity which occurs when Cy3-AB dimer is evicted from the nucleosome and the DNA undergoes productive unwrapping. Unproductive DNA unwrapping events that are not coincident with histone eviction are marked by black asterisks. (**B**) The fraction of nucleosomes showing a spontaneous increase in Cy3 signal up to five minutes after flowing different combinations of reactants into the imaging chamber. The number of traces analyzed are shown above each column. (**C**) The inverse lifetime (+/− standard deviation) of the high Cy3-H2A signal (τ in (A)) in the presence of increasing concentrations of the histone chaperone Nap1. The red line is a linear fit to the data. (**D**) The high Cy3-H2A signal intermediate state results from the evicted Cy3-AB remaining associated with the SWR1-nucleosome complex before it dissociates or is removed by Nap1.

The high Cy3-H2A signal state persists for tens of seconds (as in Fig. 2A), showing that the Cy3-AB dimer remains associated with the SWR1-nucleosome complex after its initial displacement from the histone octamer. The mean lifetime (τ) is 37.2 ± 1.9 s, after which the Cy3-AB either dissociates from the SWR1-nucleosome complex or photobleaches (fig. S5A). Interestingly, the inclusion of 100 nM Nap1, the major H2A-H2B histone chaperone of *S. cerevisiae*, reduces τ by 8-fold (fig. S5A) without altering the overall rate of histone displacement (fig. S5B). Nap1 does not affect Cy3 photobleaching (fig. S5C). The rate of Cy3-AB dissociation (1/ τ) increases linearly with Nap1 concentration (Fig. 2C), suggesting that Nap1 can competitively remove the AB dimer from SWR1 after its displacement from the nucleosome (Fig. 2D).

SWR1 exchanges both copies of AB dimers on a canonical nucleosome (*28*). Although the probability of observing two sequential histone eviction events is low due to photobleaching, we could indeed observe such events on 32 nucleosomes (Fig. 3A,B and additional examples in fig. S6A,B). On average, the first eviction event occurs 58.1 ± 4.5 s after introduction of reactants, similar to the time interval between the first and second eviction events (57.5 ± 14.6 s) (fig. S6C), confirming that histone exchange is distributive, meaning two rounds of histone exchange are carried out by separate SWR1 complexes (*15*). There is biochemical evidence suggesting that the Swc5 subunit of SWR1 may interact with the AB dimer to facilitate its eviction (*29, 30*). The regions on the AB dimer that interact with Swc5 and Nap1 partially overlap, raising the possibility that Nap1 competes with Swc5 for the same AB binding site (*29, 31*). We speculate that transfer of evicted AB from SWR1/Swc5 to Nap1 prevents nonspecific binding of free histones to DNA after exchange.

**Fig. 3.**
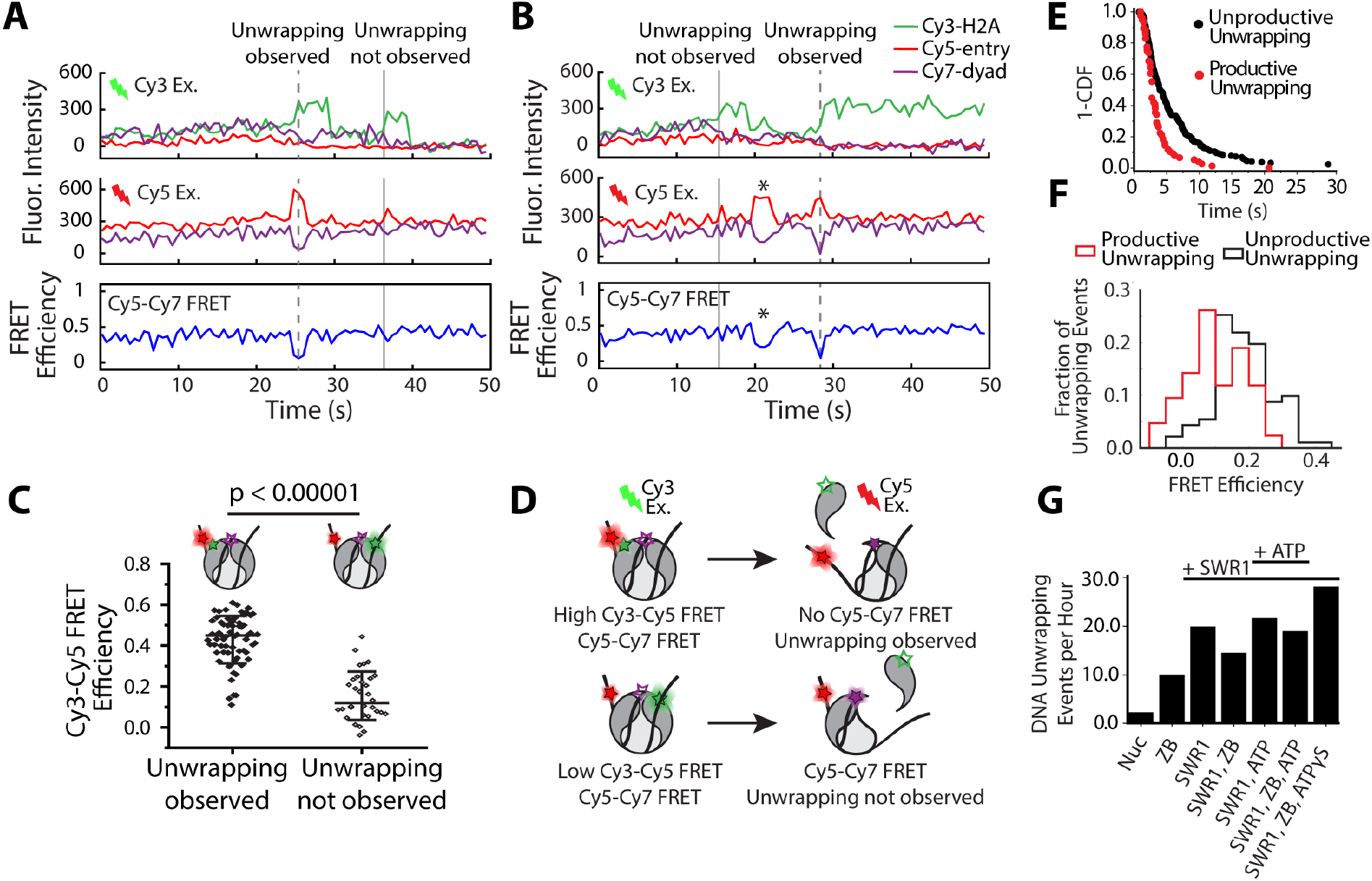
Differentiating Unwrapping and Productive DNA Unwrapping. (**A** and **B**) Some nucleosomes containing two Cy3-AB dimers exhibited two histone exchange events. In these cases, only one histone exchange event occurred simultaneously with entry DNA unwrapping and rewrapping. (**C**) DNA unwrapping events are only observed when the Cy5-proximal Cy3-AB is evicted, as is shown by the difference in Cy3-Cy5 FRET efficiencies immediately prior to histone eviction for nucleosomes containing one Cy3-AB dimer. This shows that SWR1 only unwraps DNA from one face of the nucleosome during histone exchange, as shown in (**D**). (**E**) 1-CDF of DNA unwrapping lifetimes for productive DNA unwrapping events concurrent with histone exchange, and unproductive unwrapping events that do not result in histone exchange. (**F**) The average Cy5-Cy7 FRET efficiency is lower during productive DNA unwrapping than in unproductive DNA unwrapping. (**G**) The frequency of unwrapping events detected in the presence of different combinations of reactants. The addition of SWR1 alone, without ATP and ZB dimer, can cause nucleosomes to unwrap, suggesting that SWR1 binding to nucleosomes is sufficient to cause unproductive DNA unwrapping.

The conformation of nucleosomal DNA can be monitored by measuring the FRET efficiency between Cy5 and Cy7 placed at the DNA entry and DNA dyad. Among nucleosomes that exhibit histone eviction, 45% (78/173) are accompanied by a near-simultaneous decrease of Cy5-Cy7 FRET efficiency (see Fig. 2A, Fig. 3A,B, fig. S4A,B, fig. S5D,E, fig. S6A,B), suggesting that DNA unwrapping is coincident with AB eviction. We name unwrapping events that are coincident with histone eviction productive unwrapping. Not every DNA unwrapping event is accompanied by a Cy3-H2A fluorescence increase; indeed, most unwrapping events (218/296 events on 173 nucleosomes where histone exchange is observed) do not lead to Cy3-AB dimer eviction and are called unproductive unwrapping (Fig. 2A, Fig. 3B black asterisks).

Interestingly, among nucleosomes that undergo histone eviction, 55% (95/173) display no apparent DNA unwrapping (Fig. 3A,B, fig. S4C,D, fig. S6A,B). We hypothesize that SWR1 only unwraps DNA from the side of the nucleosome where AB is evicted, leaving DNA on the opposite face of the nucleosome wrapped. Thus, if SWR1 unwraps DNA from only the unlabeled ‘exit’ side when the dimer in the exit side is evicted, the entry-dyad FRET would be blind to such unwrapping (Fig. 3D). We tested this hypothesis by measuring FRET between Cy3-H2A and Cy5 at the DNA entry site just prior to histone eviction. The FRET efficiency is indeed higher for eviction coincident with unwrapping (0.43 +/− 0.12 (N=66)) compared to eviction not coincident with unwrapping (0.15 +/− 0.12 (N=32) (Fig. 3C), showing that productive DNA unwrapping is only observed via FRET when the AB dimer proximal to the Cy5-DNA label is evicted. In further support, nucleosomes showing two sequential eviction events display a transient decrease of Cy5-Cy7 FRET solely when the evicted Cy3-AB dimer is proximal to the Cy5-DNA label (fig. S6A,B). These findings provide strong evidence that SWR1 only unwraps DNA on the same nucleosome face as the AB dimer to be displaced, leaving DNA on the opposite face unperturbed (Fig. 3D).

To elucidate the difference between productive and unproductive DNA unwrapping, we measured the lifetime of the unwrapped state and the extent of DNA displacement by calculating the average Cy5-Cy7 FRET efficiency during the unwrapping event. Productive DNA unwrapping events are 2-fold shorter (2.3 s +/− 0.2 s) than unproductive unwrapping events (4.2 s +/− 0.1 s) (Fig. 3E). In addition, the time-averaged Cy5-Cy7 FRET efficiency is significantly lower during the productively unwrapped state (0.1 +/− 0.01) than during unproductive DNA unwrapping events (0.18 +/− 0.01) (Fig. 3F). Hence, the two characteristics that distinguish productive from unproductive DNA unwrapping are a larger unwrapping amplitude and more rapid temporal recovery from the unwrapped state.

Based on the measured change in FRET efficiency, we estimate that DNA is displaced 2.2 nm from its usual path around the histone octamer during productive unwrapping (see Methods for distance calculations). In the cryo-EM structure 25 bp of DNA are unwrapped from one side of the nucleosome bound to SWR1 in complex with ZB dimer and an ATP analogue (*15*). The distance between the Cy5 and Cy7 DNA labeling sites in the SWR1-nucleosome structure is 6.5 nm-2.4 nm longer than their separation in the structure of the unbound, fully wrapped nucleosome (see Methods for distance calculation). This change is consistent with the 2.2 nm displacement that occurs in productive unwrapping events, and larger than the 1.2 nm displacement observed in unproductive unwrapping.

To determine which reaction components are responsible for unproductive DNA unwrapping, we measured the frequency of DNA unwrapping events (Fig. 3G). Nucleosomes alone show very low unwrapping frequency (1 unwrapping event in 27 minutes). Inclusion of ZB dimer increases the frequency 4.5-fold (1 event every 6 minutes), likely due to non-specific binding of histone dimer to the nucleosome (*32*). Addition of only SWR1 to nucleosomes further increases the frequency (1 event every 3 minutes). We found little correlation between unwrapping frequency and the inclusion of additional reaction components (ZB dimer, ZB dimer + ATP, ZB dimer + ATPγS or ATP alone) (Fig. 3G). The results show that SWR1 binding alone can unproductively unwrap nucleosomes.

While SWR1 binding alone is sufficient to cause unproductive DNA unwrapping, we do not know whether nucleosomal DNA remains unwrapped for the duration of the SWR1-bound state. To directly observe and correlate SWR1 binding with nucleosomal DNA unwrapping, we specifically labeled the SWR1 complex with Cy3 (Cy3-SWR1) and examined its real-time binding to labeled nucleosomes (Cy5-DNA entry, Cy7-DNA dyad) (fig. S7). We found that a small fraction of nucleosomes (51/397, 13%) show transient DNA unwrapping upon SWR1-binding in the absence of ATP and ZB dimer while the majority of SWR1-bound nucleosomes (346/397, 87%) remain stably wrapped (fig. S7A). Given that the lifetime of the Cy3-SWR1-nucleosome complex is photobleaching-limited in these real-time measurements, we measured the lifetime of the immobilized Cy3-SWR1-nucleosome complex in a different experiment by introducing Cy3-SWR1 to nucleosomes and imaging intermittently after a buffer wash to remove unbound molecules. These measurements showed that the SWR1-nucleosome complex is highly stable in these conditions and has a lifetime on the order of tens of minutes in the absence of ATP and ZB dimer (fig. S7B). However, in the presence of ATP alone or ATP and ZB dimer the lifetime of the bound state decreased to 14 minutes (fig. S7B). This suggests that ATP binding alone could affect the affinity of SWR1 for its substrate nucleosome.

Our results indicate that while SWR1 binding per se under experimental conditions is sufficient to partially destabilize the nucleosome and raise the frequency of DNA unwrapping, the lifetime of the unwrapping events (seconds) is significantly shorter than the bound lifetime of the SWR1-nucleosome complex (minutes). As such, the SWR1-bound nucleosome is predominantly normally wrapped (fig. S7C), which contrasts with other nucleosome remodelers, such as CHD1, that unwrap nearly 100% of the nucleosome population upon binding under saturating conditions (*33, 34*).

Unproductive unwrapping events due to SWR1 binding alone could result from SWR1-induced stabilization of the transient and normally short-lived intrinsic unwrapping of the outer gyre of nucleosomal DNA (*35, 36*). This interaction could be mediated by several subunits in the SWR1 complex (Swc2, Swr1, Arp6 and Swc6) which either contact DNA in cryo-EM structures, or have DNA binding regions that could bind and unwrap DNA (*15, 21, 37, 38*). Integrating our findings for productive and unproductive DNA unwrapping, we hypothesize that when all three natural substrates are engaged with SWR1 to maximally stimulate its ATPase activity (*28*), translocation of DNA by the catalytic ATPase at SHL2 opposite long linker DNA ruptures multiple DNA contacts on the AB dimer, unwrapping DNA to sufficiently destabilize the histone octamer and facilitate dimer exchange.

Because SWR1 does not slide the histone core relative to the DNA during histone exchange (*39*), the exact time of Cy3-AB dimer displacement from the octamer can be identified by the reduction of FRET efficiency between Cy3-AB and Cy7 at the DNA dyad. The Cy3-Cy7 FRET decrease occurs momentarily after the Cy5-Cy7 FRET decrease, indicating that dimer eviction is preceded by DNA unwrapping (Fig. 4A,B, additional example in fig. S8A,B), by 1.0 s on average (τ_U-E_, Fig. 4C, fig. S8D). After histone eviction, the mean time until the DNA returns to the rewrapped state is 1.4 s (τ_E-R_, fig. S8C). Because AB eviction and ZB insertion are tightly coupled in biochemical assays (*6, 28*), as well as in our single-molecule assay (Fig. 1D), we suggest that ZB is inserted within the 1.4 s window between histone eviction and DNA rewrapping (Fig. 4D).

**Fig. 4.**
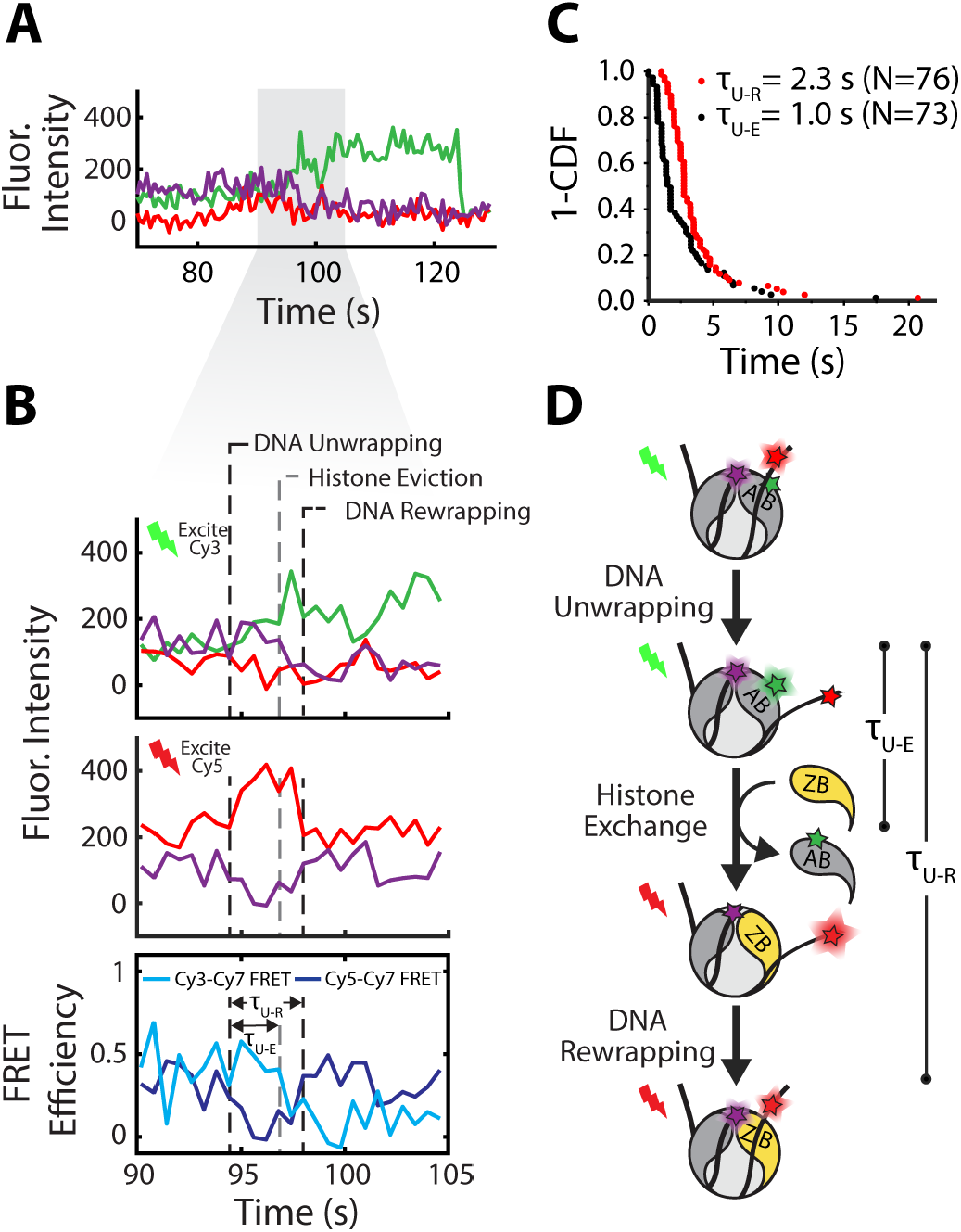
Dimer Eviction Occurs Just After DNA is Productively Unwrapped. (**A**) Fluorescence intensity trajectories of a nucleosome where histone eviction occurs after DNA unwrapping. (**B**) Expanded view of the shaded region in (A). Black dashed lines indicate DNA unwrapping and rewrapping which is observed as changes in the Cy5-Cy7 FRET efficiency. A grey dashed line indicates where histone eviction occurs, characterized by loss of Cy3-Cy7 FRET. t_U-R_ is defined as the time between DNA unwrapping and rewrapping; t_U-E_ is the time between DNA unwrapping and histone eviction. (**C**) 1-CDF of t_U-R_ and t_U-E_. (**D**) Schematic of nucleosome intermediates during productive DNA unwrapping.

These measurements revealed several new intermediates in the histone exchange reaction and allowed us to construct a detailed reaction mechanism in the context of a +1 nucleosome in *S. cerevisiae*, the most well-characterized H2A.Z-enriched nucleosome (Fig. 5). Our results support that H2A.Z enrichment on the NFR-distal side of the +1 nucleosome is caused by the mechanistic preference of SWR1 in H2A.Z deposition. This suggests that H2A.Z is likely to be enriched on the linker-distal side of other H2A.Z-containing nucleosomes, such as those next to CTCF binding sites (*40*). Once bound to its nucleosome substrate and ZB dimer, the ATPase activity of SWR1 is maximally stimulated, driving a large DNA displacement from one face of the nucleosome that initiates the exchange reaction and primes AB for removal. The 2.3 second time gap from productive DNA unwrapping to rewrapping after histone exchange shows that SWR1-mediated H2A.Z deposition is intrinsically rapid. The timescale for histone exchange is very similar to the ATP-driven nucleosome sliding at a few base pairs per second by the ISWI, SWI/SNF, CHD1 and INO80 remodelers (*34, 41–45*). This indicates that all remodelers operate on the same time scale to efficiently maintain a poised chromatin architecture at yeast promoters and other physiological locations.

**Fig. 5.**
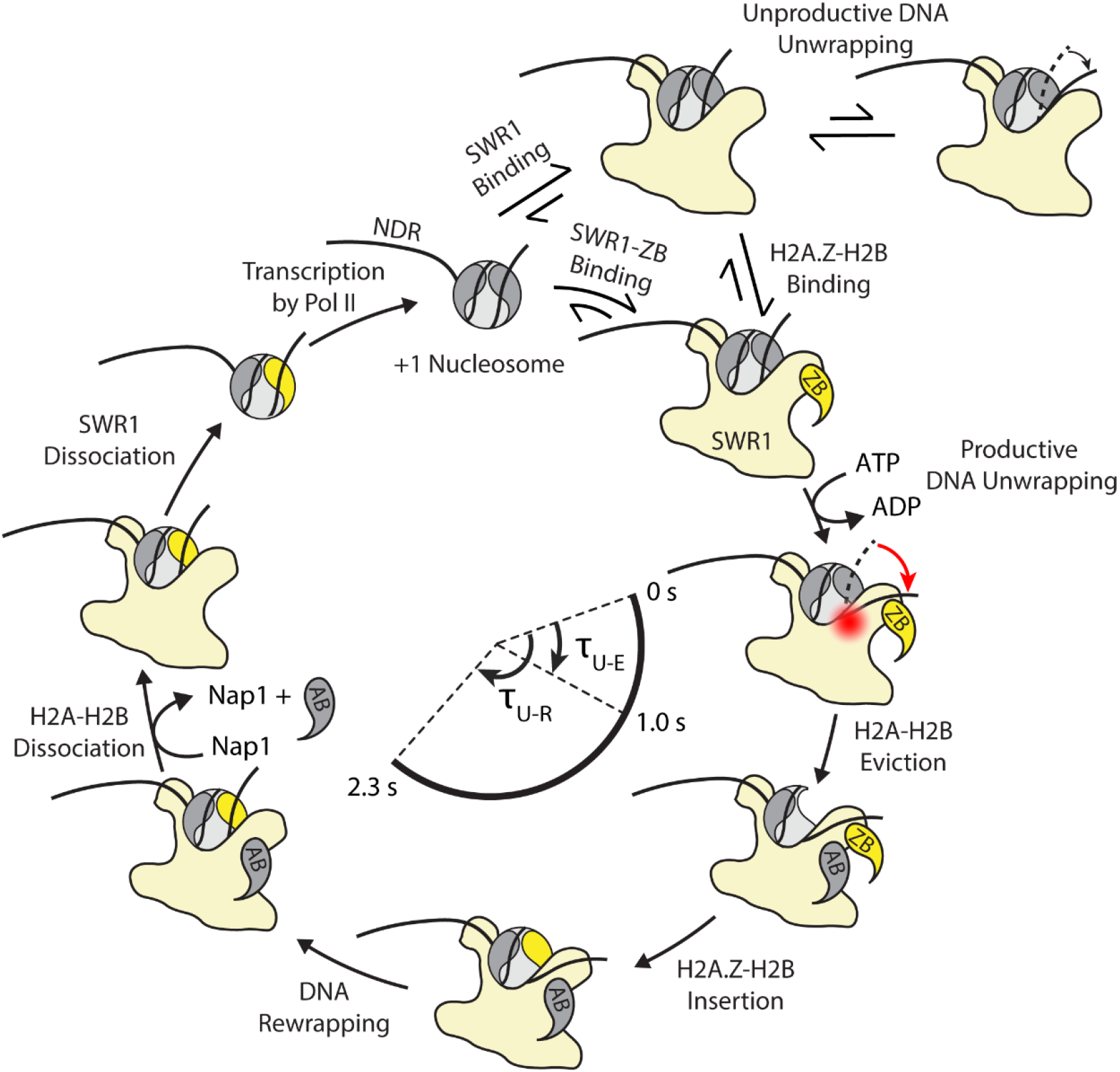
The histone exchange cycle at the +1 nucleosome in yeast. New intermediates in the histone exchange reaction cycle revealed by three-color smFRET measurements. First, SWR1 binds to the +1 nucleosome with its ATPase domain oriented distal to the NDR. Once bound, SWR1 can transiently unwrap nucleosomal DNA. These unwrapping events can either be unproductive events that do not result in histone exchange, or productive events that destabilize the histone octamer leading to AB eviction. Productive DNA unwrapping requires ATP and ZB dimer. In a productive DNA unwrapping event, the energy from ATP hydrolysis is used to unwrap DNA more extensively from the histone octamer, as shown by the red arrow, than in an unproductive DNA unwrapping event (black arrow). Next, AB is evicted an average of 1 second after the initial productive DNA unwrapping event. While not observed in our experiments, we hypothesize that the ZB dimer is inserted into the nucleosome, followed by DNA rewrapping. The average time that DNA remains productively unwrapped is 2.3 seconds. Even after eviction from the histone octamer, the AB dimer remains associated with the SWR1-nucleosome complex for tens of seconds, and its removal could be facilitated by the histone chaperone Nap1. During the last step of the exchange reaction SWR1 dissociates. The heterotypic nucleosome can go through the exchange cycle a second time for replacement of the NDR-proximal AB, or it can be disassembled via transcription elongation.

These identified reaction intermediates also put previous findings into a new context. The cryo-EM structure of SWR1 bound to the nucleosome clearly shows that a substantial amount of DNA is unwrapped upon SWR1 binding in the presence of ZB dimer and an ATP analogue, ADP-BeF (*15*). Our finding that the nucleosomal DNA remains wrapped for the majority of the time SWR1 is bound and only unwrapping productively for 2.3s during histone exchange suggests that the reported cryo-EM structure represents a transient intermediate in the exchange reaction where SWR1 is distorting DNA at SHL2 to stimulate productive unwrapping and destabilize the histone octamer. While INO80 and SWR1 share many homologous subunits and a similar architecture, the Ino80 ATPase uniquely contacts the nucleosome at SHL6 as opposed to SHL2. The difference in contact point could be the reason INO80 only unwraps a small amount of DNA from the entry site of the nucleosome (*46, 47*), similar to the unproductive DNA unwrapping caused by SWR1 binding alone. This level of unwrapping may be sufficient for INO80 to reposition histone octamers on DNA, its primary biological function.

The three-color single-nucleosome FRET assay offers a powerful approach to identify additional histone exchange reaction intermediates, elucidate the role of specific SWR1 subunits in transducing the chemical energy of ATP hydrolysis into the work of histone exchange, and assess the physical influence of DNA sequence (*24, 25, 27, 48, 49*) and the role of histone modifications in remodeling reactions (*21, 50*). More generally, multi-color single-nucleosome FRET provides a complementary approach to quantify the real-time dynamics of chromatin-based regulators and their physical effects on nucleosome architecture, composition, and inter-nucleosomal interactions.

## Materials and Methods

### Histone Expression and Purification

Wild type and mutated *S. cerevisiae* H2A-H2B, H2A.Z-H2B-3xFLAG, H2A.Z(K120C)-H2B-3xFLAG and H2A(K120C)-H2B were co-expressed polycistronically in a pST39 plasmid vector. *S. cerevisiae* H2A.Z(K120C), *S. cerevisiae* H2B, *D. melanogaste*r H3 and *D. melanogaste*r H4 were expressed as monomers in pET28 plasmid vectors. All proteins were expressed in *E. coli* using previously established protocols(*51, 52*). After expression and before purification, the cell pellets containing H2A.Z(K120C) and H2B were combined. This allowed the H2A.Z(K120C)-H2B dimer to refold during the protein purification steps detailed below.

The polycistronically co-expressed H2A-H2B, H2A.Z-H2B-3xFLAG, H2A.Z(K120C)-H2B-3xFLAG, H2A(K120C)-H2B and the combined cell pellets containing H2A.Z(K120C)-H2B, were purified in non-denaturing conditions detailed elsewhere (*52*). Briefly, cells were lysed using sonication and spun down to isolate the histone-containing insoluble fraction. The insoluble pellet was resuspended in at least 30 mL of 0.25 N HCl and placed at −20 °C for one hour at minimum. The mixture was thawed, centrifuged and the supernatant containing the solubilized histone proteins was retained. Approximately 5-10 mL of 1M pH 7.6 Tris was added to histone protein extract to neutralize the pH. The histone protein solution was then dialyzed overnight into D Buffer (10 mM HEPES at pH 7.6, 1 mM EDTA, 10% glycerol, 0.2 M NaCl and 1mM beta-mercaptoethanol (BME)). Finally, histone dimers were purified using ion exchange chromatography. Dimers were immobilized on SP Sepharose Fast Flow resin (GE Healthcare) and eluted with a gradient of 0.2 to 2 M NaCl in D buffer. Histone-containing fractions were identified using SDS-PAGE, combined, concentrated, flash frozen and stored at −80°C for further use. The H2A.Z(K120C)-H2B made by combining cell pellets was also purified further by size exclusion chromatography on an S75 column (Superdex 75 Increase 10/300 GL, GE Healthcare) before storage.

H3 and H4 monomers were purified from inclusion bodies in a previously established protocol (*51*). After extraction from inclusion bodies, the H3 and H4 monomers were purified using a mono-Q HR 5/5 column (GE healthcare) and SP ion exchange resin (GE Healthcare). Following purification, the histone containing fractions were identified using SDS-PAGE, combined, dialyzed into 18 megaohm water and lyophilized. The lyophilized histone powders were stored at room temperature for later use.

### Expression & Purification of Nap1

The recombinant yeast Nap1 was overexpressed in *E. coli* and purified by His-tag affinity purification using Ni-NTA agarose beads (Qiagen) followed by ion exchange purification using a mono-Q HR 5/5 column (GE Healthcare). Purified Nap1 was snap frozen in buffer containing 10 mM HEPES pH 7.6, 1.5 mM MgCl_2_, 0.5 mM EGTA, 0.1 mM EDTA and 10% glycerol, and stored at – 80° C.

### Expression & Purification of Swc7

The coding sequence for Swc7 (C62A) was inserted into a pET-based E. coli expression vector with an N-terminal 6xHis tag followed by an MBP tag (Addgene #29654) by ligation independent cloning (LIC). The native Swc7 sequence contains two cysteines. The C62A mutation was performed to leave one cysteine in the sequence (the third amino acid in the protein) that could be specifically labeled with Cy3 (see Protein Labeling section). For expression the plasmid was transformed into E. coli BL21-CodonPlus (DE3)-RIL competent cells (Agilent). Cells were grown in Terrific Broth media to an optical density of 1.0 (600 nm), at which point the culture temperature was shifted to 30 °C and expression was induced by addition of 0.4 mM isopropyl β-D-1-thiogalactopyranoside and cells were grown for an additional 6 h. Cells were pelleted by centrifugation and resuspended in lysis buffer (4x TBS, 0.5 mM EDTA, 2.5 mM MgCl2, 0.05% Triton X-100, cOmplete protease inhibitor cocktail (Roche), 500 U benzonase (Sigma-Aldrich)). All subsequent protein purification steps were performed at 4 °C. Cell pellets were lysed via sonication then cleared by centrifugation at 18,000 g. The clarified lysate was mixed with amylose resin (New England BioLabs) pre-equilibrated in lysis buffer and incubated for 60 min on a rocking platform. The resin was washed with wash buffer 1 (4x TBS, 1 mM EDTA, 0.025% Triton X-100, 5 mM BME), then wash buffer 2 (20 mM HEPES pH 7.6, 0.2 mM EDTA, 2 mM MgCl2, 5% glycerol, 200 mM NaCl, 1 mM DTT). The protein was eluted with wash buffer 2 supplemented with 20 mM maltose. To separate Swc7 from the MBP affinity tag, His6-TEV protease was added to the amylose resin elution at a molar ratio of 1:50 and incubated overnight at 4 °C. The TEV reaction was passed through a HisTrap HP column (GE Healthcare) as a subtractive step to remove His6-TEV protease, cleaved His6-MBP tag, and uncleaved protein. The flow-through was concentrated to 0.5 mL and applied to a Superdex 200 size-exclusion column (GE Healthcare) equilibrated in column buffer (20 mM HEPES pH 7.6, 0.2 mM EDTA, 2 mM MgCl2, 5% glycerol, 200 mM NaCl, 0.5 mM TCEP). Peak fractions were pooled and concentrated before storing at −80 °C until use.

### Protein Labeling

H2A(K120C)-H2B, H2A.Z(K120C)-H2B, H2A.Z(K120C)-H2B-3xFlag and Swc7 were all cysteine-labeled with maleimide-functionalized Cy3 dye (GE Healthcare) using previously published protocols (*53*). Briefly, H2A(K120C) and H2A.Z(K120C) were labeled in their native state, dimerized with H2B or H2B-3xFlag. H2A(K120C) and H2A.Z(K120C) were labeled in 20 mM Tris and 200 mM NaCl at pH 7, at a protein concentration of 1 – 2 mg/mL. Swc7 was labeled in 20 mM HEPES pH 7.6, 0.2 mM EDTA, 2 mM MgCl2, 5% glycerol, 200 mM NaCl and 0.5 mM TCEP at a concentration of 100 µM. All proteins were labeled with a 20-to-30-fold excess of Cy3-maleimide. Labeling proceeded for 3 hours at room temperature before the reaction was quenched with 10 mM beta-mercaptoethanol (BME). Free dye was removed from the labeled proteins by successive concentrations and dilutions using a 3 K MWCO centrifugal concentrator (Amicon Ultra-4, Millipore). The labeling efficiency was computed by measuring the concentration of the protein and Cy3 dye using their absorbances at 280 nm and 550 nm, respectively.

### Purification of Native SWR1 and Cy3-SWR1

Native SWR1 was purified from *S. cerevisiae* via a tandem affinity purification (referred to as the ASAP protocol) using a 3x FLAG tag on the SWR1 subunit and an MBP tag on Rbp1 as detailed elsewhere (*24*). To construct a specifically-labeled SWR1 complex which retained its wild type activity we chose to label the Swc7 subunit, which is non-essential for exchange activity (*54*). Creating the SWR1 complex labeled only on Swc7 was done by purifying SWR1 without Swc7 (SWR1ΔSwc7) and adding recombinant, Cy3-labeled, Swc7 into the purification at a specific point to reconstitute the full SWR1 complex containing Cy3-labeled Swc7. To generate the strain to purify SWR1(ΔSwc7), Swc7 was deleted by replacement with a natMX antibiotic resistance cassette using standard yeast transformation methods in the ASAP SWR1 yeast strain (*24*), which harbors the SWR1-3xFLAG-loxP, RVB1-MBP-loxP, and htz1Δ::kanMX alleles. After elution from the amylose resin in the normal ASAP protocol, 1 nmol of Cy3-Swc7 was added to the amylose resin elution. The proteins were incubated on ice for 30 min so that Cy3-Swc7 could bind to the SWR1(ΔSwc7) and form the specifically labeled Cy3-SWR1. A glycerol gradient after all purification steps showed that Cy3-Swc7 co-migrated with the rest of the SWR1 complex, suggesting stable incorporation. After this, the normal ASAP purification protocol was followed. The reactivity of the Cy3-SWR1 complex was identical to the wild type SWR1 complex in an EMSA assay of histone exchange activity. All SWR1 complexes used in this study were separated into small, one-use aliquots to minimize freeze-thaw cycles.

### Octamer Refolding

Histone octamers were reconstituted using a previously established protocol (*51*). All experiments utilized histone octamers containing *D. melanogaster* H3 and H4 and *S. cerevisiae* H2A and H2B. These hybrid constructs were chosen as opposed to octamers made entirely of yeast histones to increase the refolding yield. Octamers made entirely of yeast histones are well known to be less stable and obtaining a high yield of octamer can be problematic (*55*). The reactivity of the hybrid octamer used in this study is similar to octamers made entirely of *S. cerevisiae* histones. The octamer reconstitution protocol is as follows. H3 and H4 monomers were denatured in 7 M Guanidinium HCl at a concentration of 2 mg/mL for two hours at room temperature. The purified H2A-H2B or Cy3-H2A(K120C)-H2B, depending on if the octamer was to be Cy3-labeled or unlabeled, was also placed into 7 M Guanidium HCl using three successive concentrations and dilutions using a centrifugal concentrator (3 K MWCO, Amicon Ultra-4, Millipore), and diluted to 2 mg/mL. After denaturing for two hours, between 0.5-2 mg of individual histones were combined and dialyzed into 2 M NaCl (10 K MWCO, 3-12 mL Slide-A-Lyzer Dialysis Cassettes, ThermoFisher Scientific). After dialysis the histones in 2 M NaCl were concentrated to a volume of 1 mL and purified using an Superdex 200 column (GE Healthcare) on an FPLC (Agilent). Octamer containing fractions were pooled, concentrated, diluted to 1 mg/mL in 40% glycerol and stored at −20 °C for later use.

### DNA Construct Formation

The biotinylated 233 bp 43-N-43 DNA (where N represents a derivative of the Widom 601 sequence (*20*), 601-c2, and 43 represents linker DNA lengths one either side of the nucleosome positioning sequence) labeled with both Cy5 and Cy7 was formed by ligating two separate PCR-generated fragments. The first fragment was a 193 bp DNA oligo labeled specifically with both Cy5 and Cy7. The construct was made via PCR amplification of a derivative of the Widom 601 sequence using Cy5 and Cy7 labeled primers (sequences of the 601-c2 derivative and primers used in this study, as well as the exact locations of the Cy5 and Cy7 labels are found in Supplemental Tables 1 & 2). The forward primer (43-N-43 Forward Primer) also contains a DraIII restriction site. The unlabeled, amine-modified primers were purchased from Integrated DNA technologies, Inc (IDT). Prior to the PCR reaction, primers were labeled with either Cy5 or Cy7 in a 62.5 µL reaction with 160 µM of amine-modified oligo, 8 mM of NHS-Cy5 (GE Healthcare) or NHS-Sulfo-Cyanine 7 (Lumiprobe) and 200 mM of fresh NaHCO_3_. The reactants were incubated with gentle mixing at room temperature for 4 hours, then at 4 °C overnight. Free dye was removed using ethanol precipitation. The labeling efficiency was 85% for Cy5 and 84% for Cy7. Next, a 2.4 mL PCR reaction was performed using Taq polymerase (NEB) in 1X GoTaq buffer to amplify a 25-N-43 DNA, where N is the 147 bp Widom 601 sequence derivative (601c2) (*20*). The PCR product was digested with the DraIII restriction enzyme (New England BioLabs) for 4 hours at 37 °C to generate a 3-N-43 fragment with a 3 nucleotide (nt) sticky overhang. The 22 bp difference between the PCR and the digestion product allowed enough separation on a native PAGE gel to confirm the digestion was complete. The digested DNA was purified on a Mini Prep Cell (BioRad) through a 6% Native PAGE column. The 37 bp biotinylated DNA fragment with 3 nt sticky overhang was formed by annealing two single-stranded DNA oligos. The 50 µL annealing reaction contains 90 µM of the 40 nt biotinylated Oligo 1 and 110 µM of the 37 nt non-biotinylated Oligo 2 in 10 mM Tris-HCl pH 8.0 and 50 mM NaCl. The annealing reaction was performed in a thermocycler, starting with 95 °C and then slowly lowering the temperature to 25 °C over an hour. The annealed DNA fragment was ligated to the Cy5-Cy7 DNA using T4 DNA ligase (ThermoFisher Scientific) for 3 hr at 22 °C followed by heat inactivation for 10 min at 65 °C. The ligated DNA was purified by phenol-chloroform extraction and on a Mini Prep Cell. The purified DNA was concentrated to 10 µM in a 0.5 mL Amicon centrifugal filter unit with 10 kDa-cutoff (ThermoFisher Scientific) and stored at −20 °C. The 233 bp 80-N-6 and 6-N-80 DNA constructs labeled with Cy5 was generated by PCR using the appropriate Cy5-labeled primes as listed in Table S2 and S3. The PCR product was purified by phenol-chloroform extraction followed by a PAGE purification on a Mini Prep Cell.

### Nucleosome Reconstitution

Nucleosome reconstitution was carried out using salt gradient dialysis as detailed previously (*51*). Briefly, nucleosomes were reconstituted by combining the desired octamer and nucleosomal DNA together, each at a concentration of approximately 262 nM, in buffer composed of 10 mM Tris at pH 7.5, 2 M NaCl, 1 mM EDTA, 1 mM BME, 0.1 mg/mL BSA and 6.4 µg of Lambda DNA (New England Biolabs). The excess lambda DNA was added to the to eliminate non-specific sticking of histones to linker DNA and prevent any aggregation that can occur in an excess of histone proteins. The nucleosome sample was placed in a 7K MWCO dialysis button (Slide-A-Lyzer MINI dialysis unit, ThermoFisher Scientific) and dialyzed from 600 mL of high-salt buffer (10 mM Tris at pH 7.5, 2 M NaCl, 1 mM EDTA, 3.5 mM BME and 0.02% NP40) into low-salt buffer (10 mM Tris at pH 7.5, 50 mM NaCl, 1 mM EDTA, 3.5 mM BME and 0.02% NP40) overnight via gradient dialysis. After dialysis, nucleosomes were concentrated to a volume of 100 µL and purified over a 4 mL sucrose gradient (5-20%). After purification, the nucleosome-containing fractions were identified by agarose gel electrophoresis, pooled, concentrated using a 10 K MWCO centrifugal concentrator and dialyzed into low-salt nucleosome reconstitution buffer overnight. The quality of nucleosomes was assayed using a 6% native PAGE. Nucleosomes were also quantified by ethidium bromide staining. The intensity of the nucleosome band was compared to the intensity of a known quantity of DNA standards on the same 6% native PAGE. Samples were stored in the dark at 4 °C and used for approximately one month.

### EMSA Assay for SWR1 Activity

A previously established assay was used to measure the exchange activity of SWR1 in bulk (*21*). Briefly, the desired concentration of SWR1, 1 nM nucleosomes, H2A.Z-H2B-3XFLAG and 1 mM ATP were combined in reaction buffer (25 mM HEPES-KOH pH 7.6, 0.37 mM EDTA, 5% Glycerol, 0.017% NP40, 70 mM KCl, 3.6 mM MgCl_2_ and 100 μg/ml BSA). ATP was left out for control experiments as needed. After mixing, 5 µL aliquots of the master reaction mix were taken at various timepoints. The reaction in these aliquots was immediately quenched by the addition of 0.5 µL of 2 mg/mL of salmon sperm DNA and placed on ice. After all aliquots had been taken, the progress of the reaction was assayed using a 6% native PAGE. Before loading on the gel, sucrose loading buffer was added to the reaction aliquot. Samples were electrophoresed for 90 minutes at 120 V in 0.5x TBE. Gels were imaged using a Typhoon scanner (Cy3, Cy5 and Cy7 excitation wavelengths were used as needed).

The specific reactant combinations used in each exchange reaction presented in this manuscript are as follows. The 43-N-43 nucleosome labeled with Cy3, Cy5 and Cy7 was used in combination with unlabeled H2A.Z-H2B-3xFLAG (fig. S1A). The Cy5-labeled 80-N-6 and 43-N-43 nucleosomes were used in combination with Cy3-H2A.Z-H2B-3xFLAG (Fig. 1E). In these reactions 5 nM SWR1 was mixed with 1 nM nucleosomes, 10 nM Cy3-H2A.Z-H2B-3xFLAG and 1 mM ATP.

### Three-Color Single-Molecule FRET Microscope Instrumentation

Total Internal Reflection Fluorescence (TIRF) Microscopy was performed on a Nikon Eclipse Ti microscope equipped with a Nikon perfect focus system and a home-built prism-TIRF module (*53*). The system was driven by home-built software. A Nikon 60X/1.27 NA objective (CFI Plan Apo IR 60XC WI) was used. Illumination was provided by solid-state lasers (Coherent, 641 nm; Shanghai Dream Lasers Technology, 543 nm and 750 nm). Emission was collected using long-pass filters (T540LPXR UF3, T635LPXR UF3, T760LPXR UF3) and a custom laser-blocking notch filter (ZET488/543/638/750M) from Chroma. Images were recorded using an electron-multiplying charge-coupled device (EMCCD; Andor iXon 897).

### Two-Color Single-Molecule FRET Microscope Instrumentation

Single-molecule FRET data was obtained using a two-color FRET microscope built to image Cy3 and Cy5 (*53*). Cy3 was excited by a 532 nm laser (Coherent Compass 315M) and Cy5 was excited by a 638 nm laser (Cobolt 06-MLD). The microscope was equipped with a water immersion objective (Olympus NA 1.2, 60×) and an iXON Andor Technology camera. The image projected onto the camera was spectrally separated so that Cy3 and Cy5 emission from the same area of the microscope could be imaged simultaneously. A long-pass emission filter (Semrock BLP02-561R-25) and notch filter (Chroma ZET633TopNotch) were used to filter out leakage from the 532 nm and 638 nm excitation sources. The fluorescence emission was spectrally separated by a long-pass dichroic mirror (Semrock FF640-FDi01-25 × 36).

### Quartz Slide Passivation

Quartz slides and glass coverslips were passivated with polyethylene glycol (98% PEG, 2% biotin-PEG JHU Slide Production Core) using previously established methods (*53*). Once passivated, quartz slides and coverslips were assembled into flow chambers with double-sided tape and epoxy.

### Nucleosome Immobilization & Imaging Conditions

Fluorescently labeled nucleosomes were immobilized in imaging chambers assembled from PEG-passivated quartz slides. After assembly 0.2 mg/mL neutravidin in 10 mM Tris-HCl pH 8.0 and 50 mM NaCl (T50) was flowed into the channel so that neutravidin could bind the biotinylated PEG on the quartz surface. The neutravidin solution was washed out within five minutes using T50. Next, nucleosomes containing biotinylated DNA were diluted to approximately 10 pM in reaction buffer (25 mM HEPES-KOH pH 7.6, 0.37 mM EDTA, 0.35 mM EGTA, 5% Glycerol, 0.017% NP40, 70 mM KCl, 3.6 mM MgCl_2_, 100 ug/ml BSA). Nucleosomes were incubated in the channel for approximately one minute before being flushed out with five channel volumes of reaction buffer. Next, imaging buffer (reaction buffer also containing an oxygen scavenging system, 0.8% w/v dextrose, 2 mM Trolox, 1 mg/ml glucose oxidase (Sigma Aldrich) and 500 U/ml catalase (Sigma Aldrich)) was introduced into the chamber immediately prior to imaging. Nucleosomes were always imaged prior to any experiment to validate that the density of nucleosomes immobilized on the surface of the slide was appropriate, and that all fluorophores engaged in FRET as expected.

### Three-Color Imaging for FRET Histogram Construction & Photobleaching Analysis

FRET histograms for the three-color nucleosomes (fig. S1C-E) were obtained by imaging the nucleosomes for a short duration (approximately 6 s) using the Alternating Laser Excitation (ALEX) excitation scheme (fig. S1B). ALEX was implemented by exciting Cy3, Cy5 and Cy7 individually for 200 ms each. Short movies of 30 frames were taken to build FRET histograms. A total of ten short movies were taken on different areas of the slide so that enough molecules could be imaged while minimizing photobleaching artifacts. Data was analyzed as described below in the Data Analysis section.

### Two-Color Imaging for FRET Histogram Construction & Photobleaching Analysis

FRET histograms for nucleosomes containing Cy3-H2A.Z-H2B and Cy5 on nucleosomal DNA (fig. S2A-E) were obtained by imaging the nucleosomes for a short duration using direct excitation as opposed to the ALEX scheme used for three-color FRET images. Here 10 frames were taken using Cy3 excitation (100 ms exposure time) followed by 10 frames of Cy5 excitation (also 100 ms exposure time). These short movies of 20 frames were used to build FRET histograms. A total of ten short movies were taken on different areas of the slide so that enough molecules could be imaged while minimizing photobleaching artifacts. Two-color FRET efficiencies were calculated as has been described in detail elsewhere (*53*).

### Steady State Measurements of Cy3-H2A Eviction and Cy3-H2A.Z-H2B Insertion on Immobilized Nucleosomes

Single-molecule Cy3-H2A eviction and Cy3-H2A.Z insertion rate measurements (Fig. 1B-D, fig. S1I,J, fig. S2A-F) were performed by immobilizing the appropriate nucleosome in the flow chamber, injecting the reactants into the chamber and imaging at various time points in order to generate FRET histograms. For histone eviction, nucleosomes labeled with Cy5 and Cy7 on DNA and Cy3 on H2A were immobilized in the imaging channel. After immobilization and washing, the desired concentration of SWR1, unlabeled H2A.Z-H2B and ATP in reaction buffer (25 mM HEPES-KOH pH 7.6, 0.37 mM EDTA, 0.35 mM EGTA, 5% Glycerol, 0.017% NP40, 70 mM KCl, 3.6 mM MgCl_2_, 100 mg/ml BSA, 0.8% w/v dextrose, 2 mM Trolox, 1 mg/ml glucose oxidase) were flown into the channel and the nucleosomes were imaged as time progressed. As Cy3-H2A in the immobilized nucleosomes is replaced by unlabeled H2A.Z-H2B in the reaction, histone exchange should result in a loss of Cy3-Cy5 FRET spots. Therefore, the progress of the reaction was monitored by dividing the fraction of nucleosomes exhibiting Cy3-Cy5 FRET at a specific time by the fraction of nucleosomes with Cy3-Cy5 FRET before exposure to the reactants. The Cy3-Cy5 FRET efficiencies of the three-color nucleosome was obtained as detailed in the Data Analysis section of the methods.

Measurements of histone insertion were done using nucleosomes labeled with only Cy5 and Cy7 on nucleosomal DNA. Here, 10 nM SWR1, 5 nM Cy3-H2A.Z-H2B dimer and 1 mM ATP were injected into the imaging chamber to initiate the reaction. In this reaction scheme Cy3-H2A.Z-H2B is inserted into the nucleosome during the exchange reaction, resulting in the appearance of Cy3-Cy5 FRET spots. A key consideration here, however, is that the Cy3-H2A.Z-H2B non-specifically sticks to the nucleosomes, resulting in the appearance of FRET spots that are not the result of histone exchange. To eliminate the non-specific binding of Cy3-H2A.Z-H2B to nucleosomes, the reaction was allowed to proceed for the desired length of time before the imaging chamber was washed quickly and sequentially with several volumes of three different reaction buffers. Wash buffer one was reaction buffer containing .04 mg/mL salmon sperm DNA, wash buffer two was reaction buffer containing 400 mM NaCl and wash buffer three was normal imaging buffer. These washes successfully removed almost all non-specifically bound Cy3-H2A.Z-H2B without disrupting the integrity of the nucleosomes. Therefore, all Cy3-Cy5 FRET spots were the result of true histone exchange where the Cy3-H2A.Z-H2B is integral to the histone octamer. After washing the nucleosomes were imaged to construct three-color FRET histograms as described in the data analysis section of the manuscript. In these experiments, the progress of the reaction was obtained by dividing the number of Cy3-Cy5 FRET spots at a specific time point by the total number of spots with Cy5 to obtain the fraction of nucleosomes that contain Cy3-H2A.Z.

### Imaging Histone Exchange In Real-Time

For real-time measurements of the histone exchange reaction, 50 µL of reactants containing 10 nM SWR1, 20 nM H2A.Z-H2B and 1 mM ATP (unless stated otherwise) was injected at the start of the recording (0 s) using a syringe pump at 1 mL/min flow rate. Long movies were recorded using the ALEX excitation scheme for 1500 frames total using a 200 ms excitation time (a total length of 5 min). Movies were analyzed as described in the Data Analysis section.

### Pull Down Experiments For Asymmetric H2A.Z-H2B Insertion

Bias in ZB insertion was assayed for asymmetrically positioned nucleosomes using a pulldown assay. Here, biotin-80-N-6 or 6-N-80-biotin nucleosomes labeled with Cy5 on the short linker DNA and Cy3-H2A.Z-H2B-3xFLAG were used as the substrates for the exchange reaction. The substrate concentrations were 1 nM nucleosomes, 10 nM SWR1, 10 nM Cy3-H2A.Z-H2B-3XFLAG and 1 mM ATP. The reaction was performed in a test tube and allowed to proceed for the desired amount of time before a 5 µL aliquot was taken and the reaction was quenched by the addition of 0.5 µL of 2 mg/mL salmon sperm DNA and placed on ice. After the final reaction time point was quenched, the nucleosomes were immediately imaged on the two-color FRET microscope described earlier in the methods. This imaging was done by diluting the nucleosomes to concentration of 50 pM and immobilizing them in imaging chambers. Nucleosomes were imaged using the direct excitation scheme described in the two-color FRET imaging section of the methods to generate FRET histograms. Cy3-H2A.Z-H2B incorporation into either side of the nucleosome was assayed by measuring the rate of growth of the two FRET peaks shown in Fig. 1G and fig. S3D.

The on-slide reaction where Cy3-H2A.Z-H2B was inserted into 43-N-43 nucleosomes labeled only with Cy5 and Cy7 on DNA was analyzed to detect a bias in ZB insertion for symmetrically positioned nucleosomes (fig. S2A-E). Here, the Cy3-Cy5 FRET histograms were analyzed at various time points in order to quantify the relative amounts of Cy3-H2A.Z-H2B inserted distal or proximal to the Cy5 on DNA. The 5- and 10-min time point Cy3-Cy5 FRET histograms were fit to two gaussian peaks (0.23 and 0.55 FRET, Cy5-distal and Cy5-proximal ZB dimer), while the later time points were fit with three gaussians to account for the 0.40 FRET peak belonging to nucleosomes with two Cy3-H2AZ-H2B dimers. The gaussian fits were used to obtain the normalized fraction of nucleosomes containing proximal, distal or two Cy3-H2A.Z-H2B molecules.

### Cy3-SWR1 Colocalization& Off-Rate Measurements

To determine whether SWR1 binding unwraps nucleosomal DNA we co-localized SWR1 binding with a nucleosome made using DNA labeled with Cy5 and Cy7 on the same DNA edge and dyad sites for the real-time reaction measurements (fig. S7A). These measurements were done by flowing in 5 nM Cy3-SWR1 into the imaging chamber containing immobilized nucleosomes immediately after starting to image using the standard ALEX excitation scheme. SWR1 binding to nucleosomes was detected by a sharp increase in Cy3 signal in spots that had both Cy5 and Cy7. As the ALEX imaging scheme was employed, it was possible to monitor DNA dynamics simultaneously with Cy3-SWR1 binding by observing Cy5-Cy7 FRET. Cy5-Cy7 FRET efficiencies were calculated as described in the data analysis section. SWR1 off rate measurements (fig. S7B) were made by injecting 5 nM Cy3-SWR1 into the imaging chamber and incubating for five minutes. After imaging to obtain the initial fraction of Cy3-SWR1-bound nucleosomes, unbound SWR1 was then washed out of the chamber by a 100 µL wash of imaging buffer alone, or imaging buffer containing 1 mM ATP or 1 mM ATP and 10 nM ZB dimer. The sample was then imaged over time and the number of Cy3-SWR1-colocalized nucleosomes were counted.

### Three-color FRET Data Analysis

Raw movies were processed into raw fluorescence intensity time traces using custom-written IDT scripts. The raw traces were corrected for excitation bleed through, chromatic sensitivity (gamma factor) and direct excitation (*56*). FRET efficiencies between every donor-acceptor pair were calculated by solving a system of equations that describe the relationship between the detected signal from each fluorophore and the FRET efficiencies between each FRET pair (*56*). The equations used to correct for bleed through are as follows,

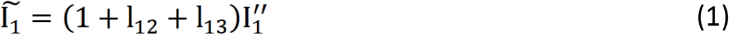

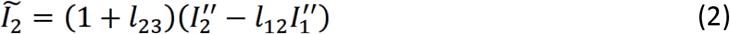

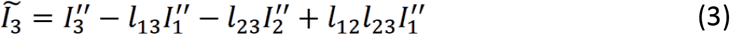

where 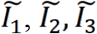 are the sum of fluorescence intensities of Cy3, Cy5 and Cy7 detected in all three channels; 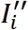 is the detected intensity of the *i*-th dye; *l_ij_* is the bleedthrough parameter of the *i*-th dye into the *j*-th channel, in units of 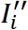. In our system, the bleed through parameters were determined as 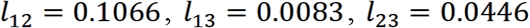 (*56*). To correct for differences in quantum yields of the fluorophores and detection efficiencies of different emission wavelengths, we determined gamma factors γ for each dye and used the corrected intensities *I_i_* for FRET efficiency calculations:

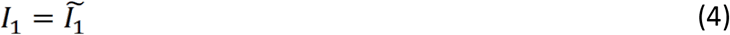

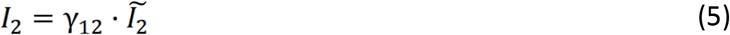

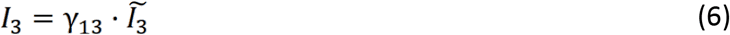

Gamma factors were determined to be 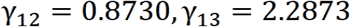 (*56*). The above corrections were applied to Cy3, Cy5 and Cy7 intensities under 543 nm (Green) or 641 nm (Red) excitation. In addition, Cy7 intensity upon 641 nm excitation is corrected for direct excitation to obtain the Cy7 intensity only from FRET with Cy5:

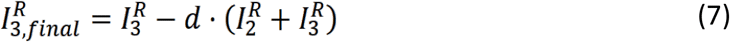

where *d* was determined to be 0.12 (*56*).

After the above corrections, FRET efficiencies between the three donor-acceptor pairs were calculated as follows:

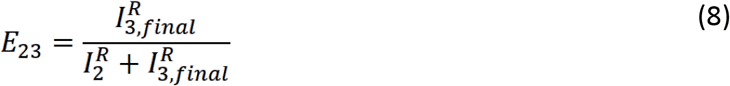

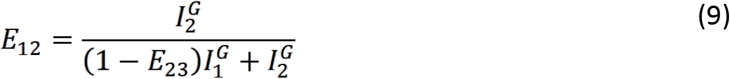

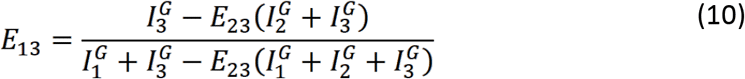

where *E*_23_, *E*_12_, *E*_13_ refer to Cy5-Cy7, Cy3-Cy5 and Cy3-Cy7 FRET efficiencies; 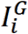 is the corrected intensity of the *i*-th dye upon green (543 nm) laser excitation; 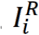 is the corrected intensity of the *i*-th dye upon red (641 nm) laser excitation (*56*).

### Dwell Time Measurements

Dwell times were collected by manual inspection using custom MATLAB scripts. ‘High Cy3 signal time’ or τ is the time from an increase in Cy3 signal due to Cy3-AB eviction to the complete loss of Cy3 signal due to dissociation of Cy3-AB from the SWR1-nucleosome complex or Cy3 photobleaching. Cy3 signal increase that can be fully explained by a decrease in Cy5-Cy7 FRET (DNA unwrapping) is excluded from this analysis. As depicted in Fig. 4C,D and fig. S6C,D, three categories of dwell times were collected for trajectories that show productive DNA unwrapping. A productive DNA unwrapping event is defined as a window of low Cy5-Cy7 FRET efficiency during which Cy3-Cy7 and Cy3-Cy5 FRET is lost due to Cy3-AB eviction. For each trace that shows productive DNA unwrapping, τ_U-E_ is the time from when Cy5-Cy7 FRET starts to decrease (DNA unwraps) to the time when Cy3-Cy7 FRET starts to decrease (H2A-H2B is evicted). τ_E-R_ is the time from when Cy3-Cy7 starts to decrease (H2A-H2B is evicted) to the time when Cy5-Cy7 FRET is completely recovered (DNA rewraps). τ_U-R_ is duration of DNA unwrapping, or the sum of τ_U-E_ and τ_E-R_. An unproductive DNA unwrapping event is defined as a window of low Cy5-Cy7-DNA FRET efficiency during which Cy3-Cy7 FRET and Cy3-Cy5 FRET (FRET between H2A and either acceptor on DNA) does not change (H2A-H2B remains in the nucleosome).

### Analysis of Cy3-Cy5 FRET Efficiency Prior to Histone Eviction

To calculate the FRET efficiency prior to histone eviction as shown in Fig. 3C, a custom MATLAB script was used. Cy3-Cy5 FRET efficiencies during a 5-10 s window before H2A-H2B eviction were averaged for each trajectory analyzed.

### Calculation of DNA Unwrapping Frequency, Unwrapped Lifetime and DNA Displacement During Unwrapping

The DNA unwrapping frequency shown in Fig. 3G was calculated by dividing the number of DNA unwrapping events by the length of the trajectory when both Cy5 and Cy7 were present to account for the effect of photobleaching. The lifetime of each productive and unproductive unwrapping events (Fig. 3E) was extracted by fitting 1-CDF of the dwell times to a single-exponential decay function. For Cy5-Cy7 FRET efficiency during productive or unproductive DNA unwrapping in Fig. 3F, the Cy5-Cy7 FRET efficiencies during the window of low Cy5-Cy7 FRET were averaged for each trajectory analyzed.

### Calculating Distances From FRET Efficiency & Comparison With Cryo-EM Structures

Distances were calculated from FRET efficiency using Eq. 11, where *E* is the measured FRET efficiency and *R_0_* is the Forster radius. As *R_0_* values for FRET pairs containing the Sulfo-Cyanine 7 dye used in this study are not published, *R_0_* values were calculated for the Cy3-Cy7 and Cy5-Cy7 FRET pairs. The Forster radius for each FRET pair was calculated using previously published methods (*57*). Briefly, the Forster radius was calculated using Eq. 12 where *N_A_* is Avogadro’s constant, *κ*^2^ is the orientation factor (taken to be 2/3), *Q_D_* is the quantum yield of the donor, *n* is the refractive index of solution (taken to be 1.33), and *J* is the spectral overlap integral. The quantum yield used for Cy3 and Cy5 were 0.16 and 0.27, respectively, as obtained from the literature (*56*). The spectral overlap integral was calculated using Eq. 13 where *f_D_(λ)* is the area-normalized donor emission spectrum, *λ* is the wavelength and *ε_A_(λ)* is the acceptor extinction coefficient spectrum. *ε_A_(λ)* was calculated by multiplying the height-normalized absorption spectra of Cy7 by its extinction coefficient, which is 240,600 mol^-1^ cm^-1^ (obtained from Lumiprobe). The calculated Forster radius for the Cy3-Cy7 FRET pair was 4.0 nm, while the Forster radius of the Cy5-Cy7 FRET pair was 6.5 nm.

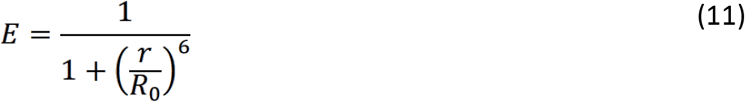

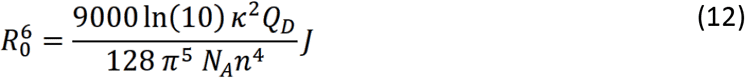

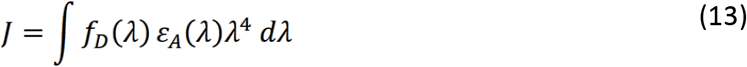

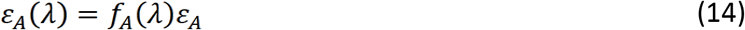

FRET Efficiencies were converted into distances using Eq. 11.

The average Cy5-Cy7 FRET efficiencies were used to measure the extent of DNA displacement during productive DNA unwrapping events. DNA displacement was calculated by finding the difference in distance between Cy5 and Cy7 in a fully wrapped nucleosome (0.35 FRET Efficiency- 7.2 nm) and the distance of DNA during a productive DNA unwrapping event (0.10 FRET Efficiency- 9.4 nm). As such, the distance between the DNA edge (Cy5) and DNA dyad increases 2.2 nm in productive DNA unwrapping events. This is significantly larger than the 1.2 nm change in distance seen in unproductive unwrapping events obtained from performing a similar calculation.

We compared the DNA displacement in productive DNA unwrapping events (2.2 nm) to the displacement of DNA observed in the SWR1-Nucleosome structure. Chimera was used to visualize the structure of the SWR1-bound nucleosome (MODEL # 6GEJ,(*15*)) and to measure the distance between the equivalent Cy7- and Cy5-labeled DNA bases in the 43-N-43 construct used in this study. In this structure, the distance between the equivalent sites was 6.5 nm. This was compared to the distance between the equivalent Cy5 and Cy7 labeling sites in the canonically wrapped nucleosome (MODEL # 1AOI,(*2*))(*32*). Note that this structure contains 147 bp of nucleosomal DNA without extra linker DNA. The distance between the first base of entry DNA at the edge of the nucleosome and the equivalent nucleotide that has been Cy7 labeled (at the DNA gyre) is 3.6 nm. The distance between the last base pair of entry DNA and the Cy5-labeled nucleotide was estimated. Assuming 0.34 nm per bp, the Cy5-labeled oligo would be 2 nm (6 bp) farther from the first DNA bp in the structure of the canonically wrapped nucleosome. Finally, we used the Pythagorean theorem to triangulate the distance between the estimated position of the Cy5-labeled nucleotide and the Cy7-labeled nucleotide. This calculated distance came out to be 4.1 nm. Therefore, the displacement induced upon SWR1 binding as observed in the structure is 2.4 nm.

## Acknowledgments

The authors thank all members of the Ha and Wu labs for support and suggestions, especially Vu Q. Nguyen and Nathan Jones for valuable comments on the manuscript. We thank Zhengjian Zhang, Andrey Revyakin, and Robert Tjian for an exploratory experiment performed at the HHMI Janelia Research Campus.

## Funding

NCI Intramural Research Program (CW)

National Institutes of Health grant R01 GM125831 (CW)

National Institutes of Health grant R35 GM122569 (TH)

National Institutes of Health Postdoctoral Training Fellowship F32 GM128299 (MFP)

National Institutes of Health Postdoctoral Training Fellowship F32 GM 133151 (R.K.L)

Howard Hughes Medical Institute (TH)

Bloomberg Distinguished Professorships (TH, CW)

## Author contributions

C.W. initially conceived the project. M.F.P and X.A.F designed the experiments. M.F.P. demonstrated initial feasibility of observing histone exchange on TIRF. M.F.P, X.A.F and J.Z. performed steady-state measurements of histone eviction and insertion. X.A.F. performed real-time histone exchange experiments. M.F.P performed experiments investigating exchange bias. M.F.P., X.A.F., F.W., and J.Y. prepared recombinant proteins and nucleoprotein complexes. M.H.J. constructed the microscope used for three-color imaging. A.R., Q.L., R.K.L., G.P. and S.L. purified and reconstituted native SWR1 complexes. X.A.F, M.F.P and M.H.J. wrote software for data analysis. M.F.P. and X.A.F. analyzed and interpreted all data. M.F.P, X.A.F., T.H. and C.W. wrote the manuscript. T.H. and C.W. supervised the project.

## Data and materials availability

Plasmids generated in this study are available upon request. Scripts used for data analysis are available at https://github.com/ashleefeng/singlemolecules/tree/master/smfret3color.

## Supplementary Materials

Supplementary Text

Figs. S1 to S8

Tables S1 to S3

## Supplementary Text

**Fig. S1.**
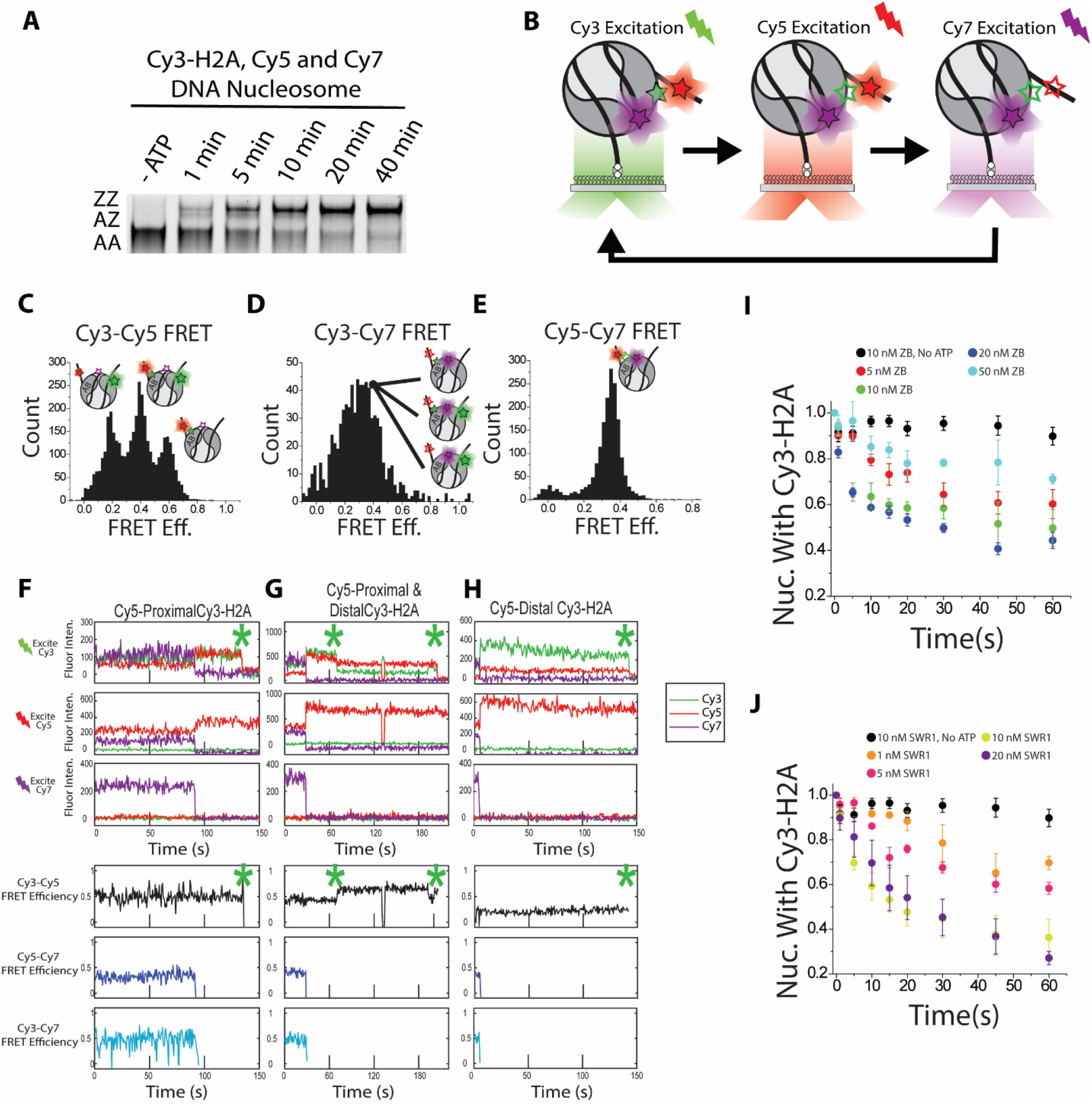
The 3-color nucleosome is a viable substrate for histone exchange. (**A**) A native PAGE assay shows the nucleosomes labeled with Cy3, Cy5 and Cy7 can undergo robust histone exchange in bulk. The gel shows time-course exchange products for the 3-color nucleosome containing Cy3-H2A(K120C), Cy5-DNA (+6 bp from entry site) and Cy7-DNA (at the DNA gyre). The reaction conditions were 1 nM nucleosome, 5 nM SWR1, 10 nM ZB-3xFLAG and 1 mM ATP. The 3xFLAG tag allows separation of AZ and ZZ nucleosomes on the gel as indicated to the left. The reaction rate is similar to unlabeled nucleosomes suggesting the Cy7 fluorophore near the DNA gyre did not disrupt histone exchange activity (*21*). (**B**), Nucleosomes were imaged using an alternating excitation (ALEX) scheme (*56*). 3-color imaging was performed by alternating between Cy3, Cy5 and Cy7 excitation for every frame during image acquisition. Upon Cy3 excitation we expect to see emission from all three fluorophores due to FRET between Cy3-H2A and Cy5 on the DNA edge, as well as FRET between Cy3-H2A and Cy7 on the DNA gyre. Upon Cy5 excitation we expect to see signal from Cy5 and Cy7 due to FRET between both Cy5 and Cy7 on DNA. Upon Cy7 excitation we only expect to see Cy7 signal which allows us to distinguish actual FRET changes from Cy7 photobleaching or blinking. (**C-E**), FRET Efficiency histograms for nucleosomes labeled with Cy3 on H2A(K120C), and Cy5 and Cy7 on the nucleosomal DNA. **C**, The Cy3-Cy5 FRET histogram showing low (0.2), middle (0.4) and high (0.58) FRET populations. The 0.2 FRET population represents nucleosomes with one Cy3-H2A distal to Cy5 on DNA. The 0.58 FRET population represents nucleosomes with one Cy3-H2A that is proximal to the Cy5 on DNA. The middle 0.40 FRET peak corresponds to nucleosomes with two Cy3-H2A histones. Cartoons depicting the labeling configuration that belong to each FRET efficiency are placed above the corresponding peak. Assignments were confirmed using photobleaching traces as shown in **F-H**. (**D**) The Cy3-Cy7 FRET efficiency histogram. Note that the three species observable in the Cy3-Cy5 FRET histogram in fig. S1C are not resolved in this FRET histogram which shows a single peak at 0.30. This could be due to the lower Förster radius (4.0 nm) of the Cy3-Cy7 FRET pair compared to the Cy3-Cy5 FRET pair (5.5 nm). (**E**) The Cy5-Cy7 FRET efficiency histogram showing a single peak at 0.38 FRET. The small peak at 0.0 FRET efficiency is assigned to the small amount of free DNA that was present in the nucleosome sample. (**F-H**), Representative time traces for nucleosomes with (**F**) one Cy3-H2A proximal to Cy5 on DNA, (**G**) two Cy3-H2A and (**H**) one Cy3-H2A distal to Cy5 on DNA. The top three plots in (**F-H**) show the intensity of Cy3, Cy5 and Cy7 when each fluorophore was excited directly using the ALEX scheme shown in fig. S1B. The calculated FRET efficiencies for all three FRET pairs are shown in the bottom three plots. Cy3 photobleaching is marked by a green asterisk. In (**G**) the Cy5-distal Cy3-H2A photobleached first as shown by the transition in Cy3-Cy5 FRET from 0.40 to 0.60 FRET efficiency after the first Cy3-bleaching event. The average Cy3-Cy5 FRET efficiency over time for these nucleosomes before Cy3-photobleaching is in agreement with the FRET histograms shown in fig. S1C-E, confirming our assignments of the 0.20, 0.58 and 0.40 FRET efficiencies to nucleosomes with one Cy3-H2A that is distal (0.20 FRET) or proximal (0.58 FRET) to the Cy5 on DNA, and that the 0.40 FRET efficiency peak represents a nucleosome containing two Cy3-H2A-H2B dimers. (**I, J**) Histone exchange rates measured by monitoring histone eviction via the loss of Cy3-H2A colocalized with Cy5 on the nucleosomal DNA as shown in Fig. 1B. (**I**) Measurements of the Cy3-AB eviction in the presence of 20 nM SWR1 and 1 mM ATP with various ZB concentrations. The histone eviction rate was fastest at 20 nM ZB. At higher concentrations, the reaction rate decreased, likely due to ZB non-specifically binding to the nucleosome and inhibiting SWR1 binding (*32, 45*). (**J**) Measurements of the Cy3-AB eviction in the presence of 20 nM ZB, 1 mM ATP and various SWR1 concentrations. The reaction rate increases with SWR1 concentration up to 10 nM SWR1. All experiments in **I** and **J** are averages of three technical replicates and the error bars are +/− standard deviation.

**Fig. S2.**
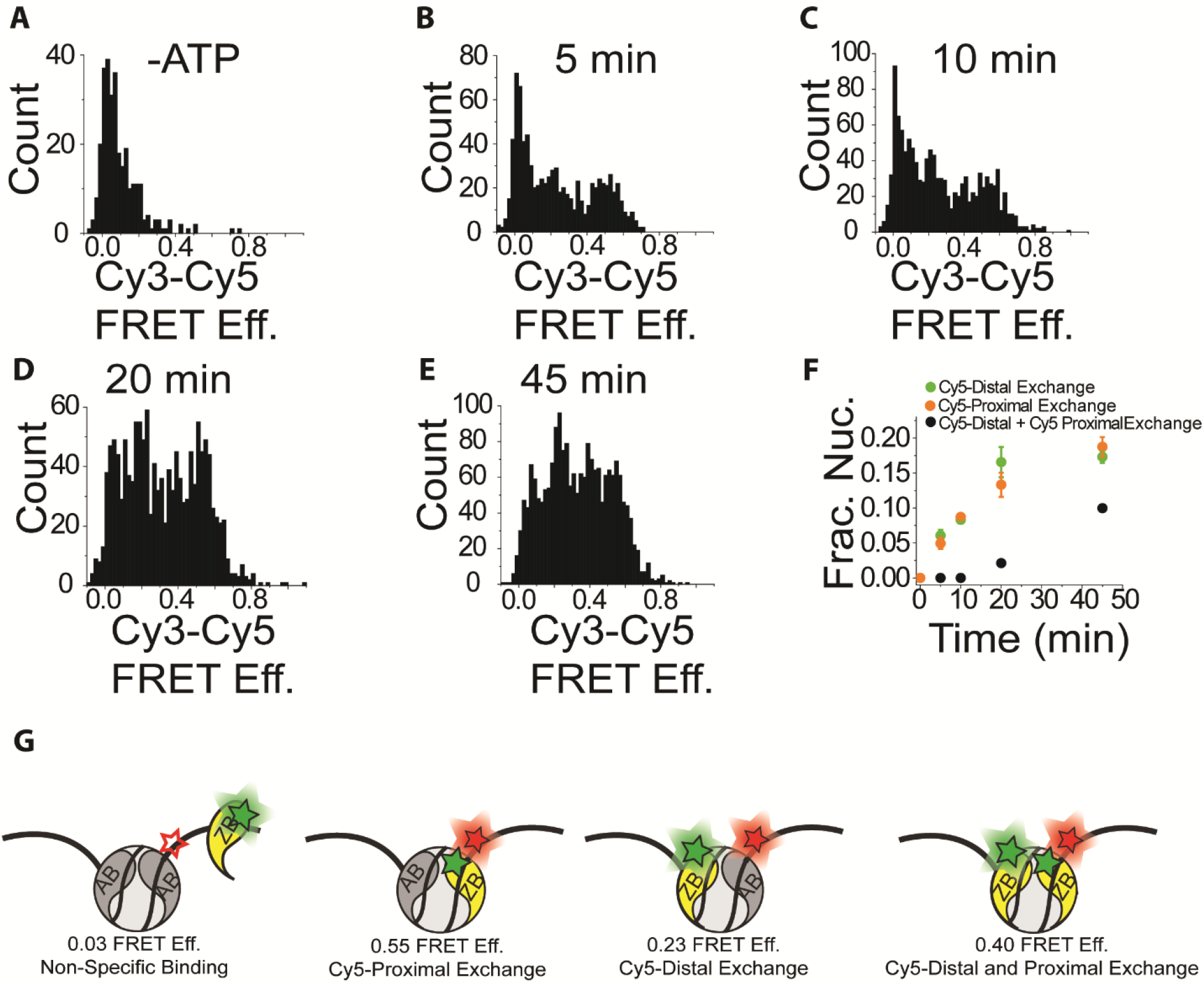
Cy3-Cy5 FRET Can Distinguish Cy5-Proximal and Distal Cy3-ZB. FRET efficiency histograms of nucleosomes exposed to SWR1, Cy3-ZB and ATP, pulled down onto quartz slides and imaged. The nucleosomes are labeled with Cy5 and Cy7 on the nucleosomal DNA. True Cy3-H2A.Z incorporation can be distinguished from non-specific binding by the Cy3-Cy5 FRET efficiency. (**A**) When ATP was left out, only a single peak was observed at 3% FRET efficiency, likely due to non-specific binding. (**B-E**) In the presence of ATP and SWR1, Cy3-ZB insertion into the nucleosome is detected as either low, medium or high FRET peaks. The 0.23 Cy3-Cy5 FRET peak corresponds to nucleosomes with Cy5-distal Cy3-ZB. The 0.55 FRET peak corresponds to Cy5-proximal Cy3-ZB and the 0.40 FRET efficiency peak corresponds to nucleosomes with two Cy3-ZB dimers. These histograms were fit to three Gaussian peaks to account for 0.23, 0.40 and 0.55 FRET populations. (**F**) The normalized growth of each peak over time, which shows that the Cy5-distal and Cy5-proximal Cy3-ZB populations grow at the same rate. As discussed in the manuscript this shows that there is no bias for histone exchange on one particular face of this symmetrically positioned nucleosome. (**G**) A cartoon depiction of the four nucleosome species corresponding to zero, low, high and mid FRET depending on if Cy3-ZB is non-specifically bound or where Cy3-ZB is inserted into the nucleosome.

**Fig. S3.**
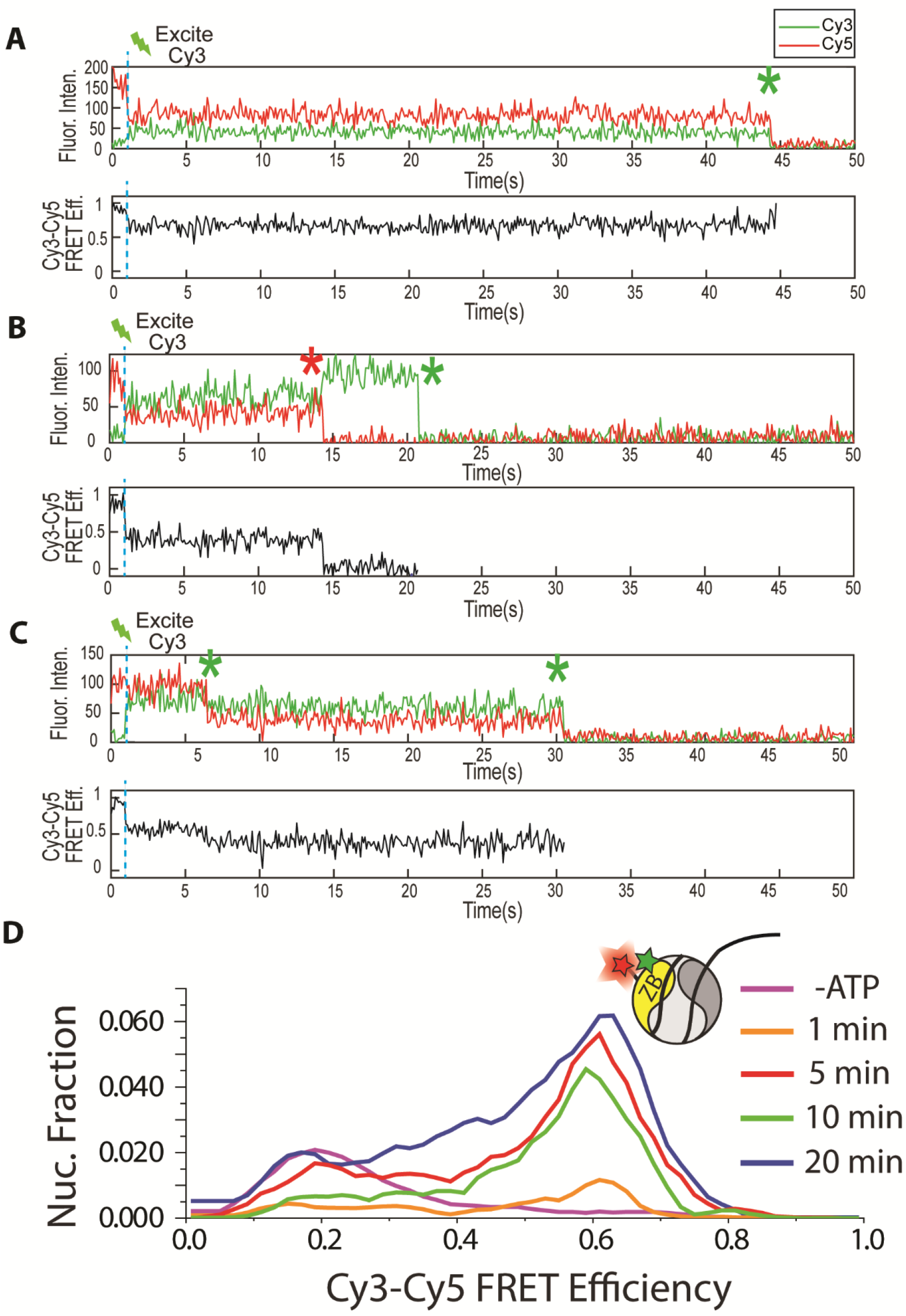
Example Traces & The Linker Distal Bias Remains for 6N80 nucleosomes. (**A-C**) Example time traces for the 80-N-6 nucleosomes pulled down after participating in a bulk histone exchange reaction where unlabeled AB is replaced by Cy3-ZB. The nucleosome is labeled with Cy5 on the nucleosomal DNA. The first second of each trace was taken under Cy5 excitation while the remainder of the trace (starting at the dashed blue line) was taken under continuous Cy3 excitation. The first second of the trace was used to identify nucleosomes labeled with a single Cy5 fluorophore, while the remainder of the trajectory was used to determine Cy3-Cy5 FRET before and after Cy3 photobleaching. The Cy3 photobleaching steps are marked by green asterisks and Cy5 photobleaching steps, if observed, by red asterisks. (**A**) A representative high-FRET trace where Cy3-ZB is proximal to Cy5 on DNA. (**B**) A representative low/mid-FRET trace where Cy3-ZB is distal to Cy5 on DNA. **(C**) A representative trace for a nucleosome that contains two Cy3-ZB. The FRET efficiency changes from mid-FRET to mid/low-FRET upon the first Cy3 photobleaching event marked by the green asterisk. This FRET change is characteristic of the Cy5-proximal Cy3-ZB photobleaching first, leaving only Cy5-distal Cy3-ZB (same FRET efficiency as in B). **(D**) The Cy3-Cy5 FRET efficiency histogram for Cy5-6-N-80 nucleosomes after exchanging with Cy3-ZB for the specified amounts of time. Nucleosomes were pulled down on to quartz slides and imaged. The FRET histograms show that just like the 80-N-6 nucleosome in Fig. 1G, Cy3-ZB is exchanged with the linker-distal dimer (high FRET population) more quickly than the linker-proximal dimer (low FRET population). That 80-N-6 and 6-N-80 nucleosomes both behave similarly show that histone exchange in this case is biased to occur on the face of the nucleosome distal to long linker DNA is on as opposed to the physical properties of the 601-nucleosome positioning sequence used in these experiments.

**Fig. S4.**
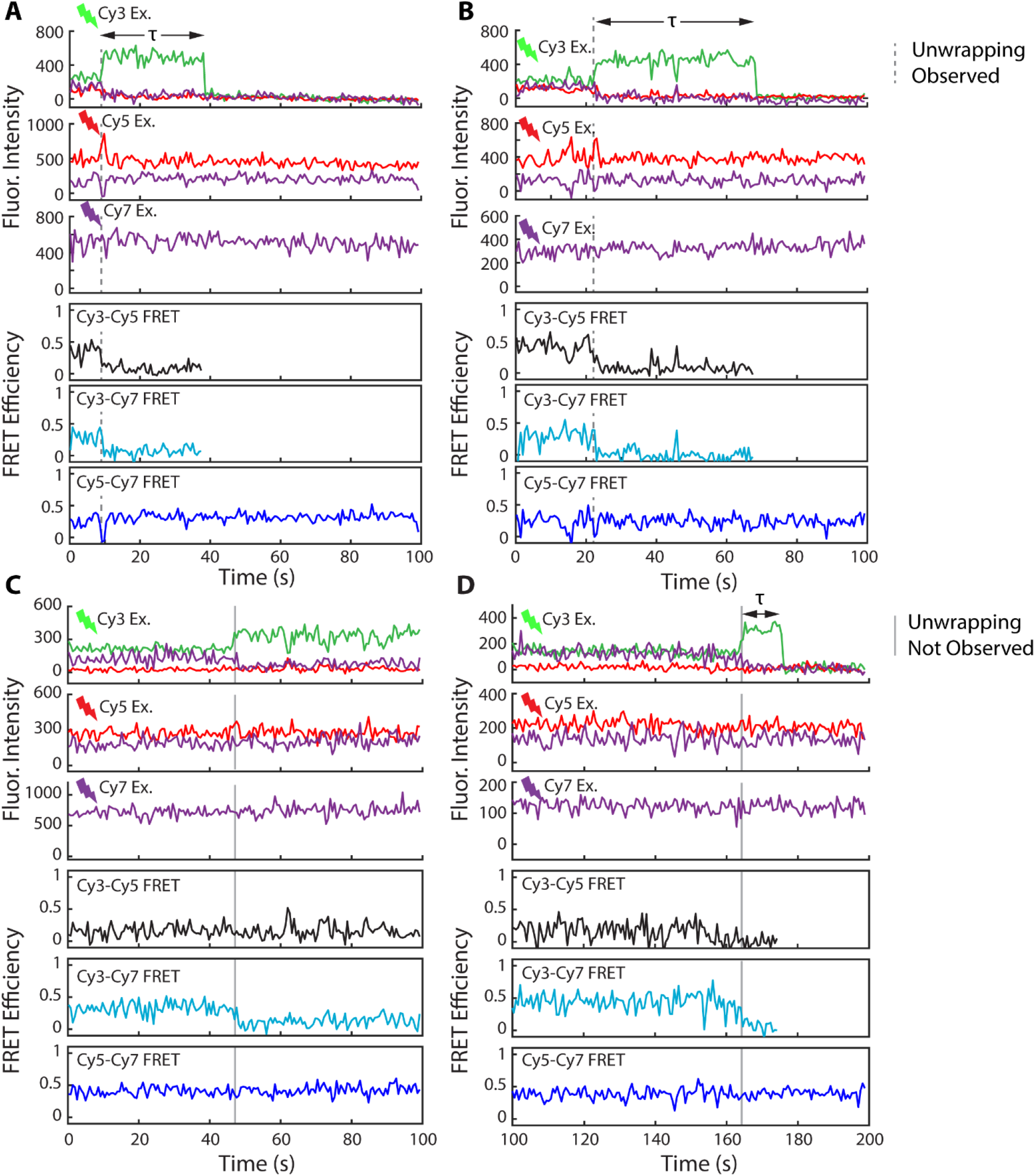
Additional single-molecule time traces showing Cy3-H2A intensity increase caused by histone eviction. (**A,B**) Histone eviction marked by Cy3-signal increase (grey dashed line) with coincident loss of Cy5-Cy7 FRET. These time traces are characterized by relatively high Cy3-Cy5 FRET prior to eviction. (**C,D**) Histone eviction (grey solid line) with no change in Cy5-Cy7 FRET at the moment of Cy3 signal increase, which marks histone eviction. These time traces are characterized by lower Cy3-Cy5 FRET prior to eviction.

**Fig. S5.**
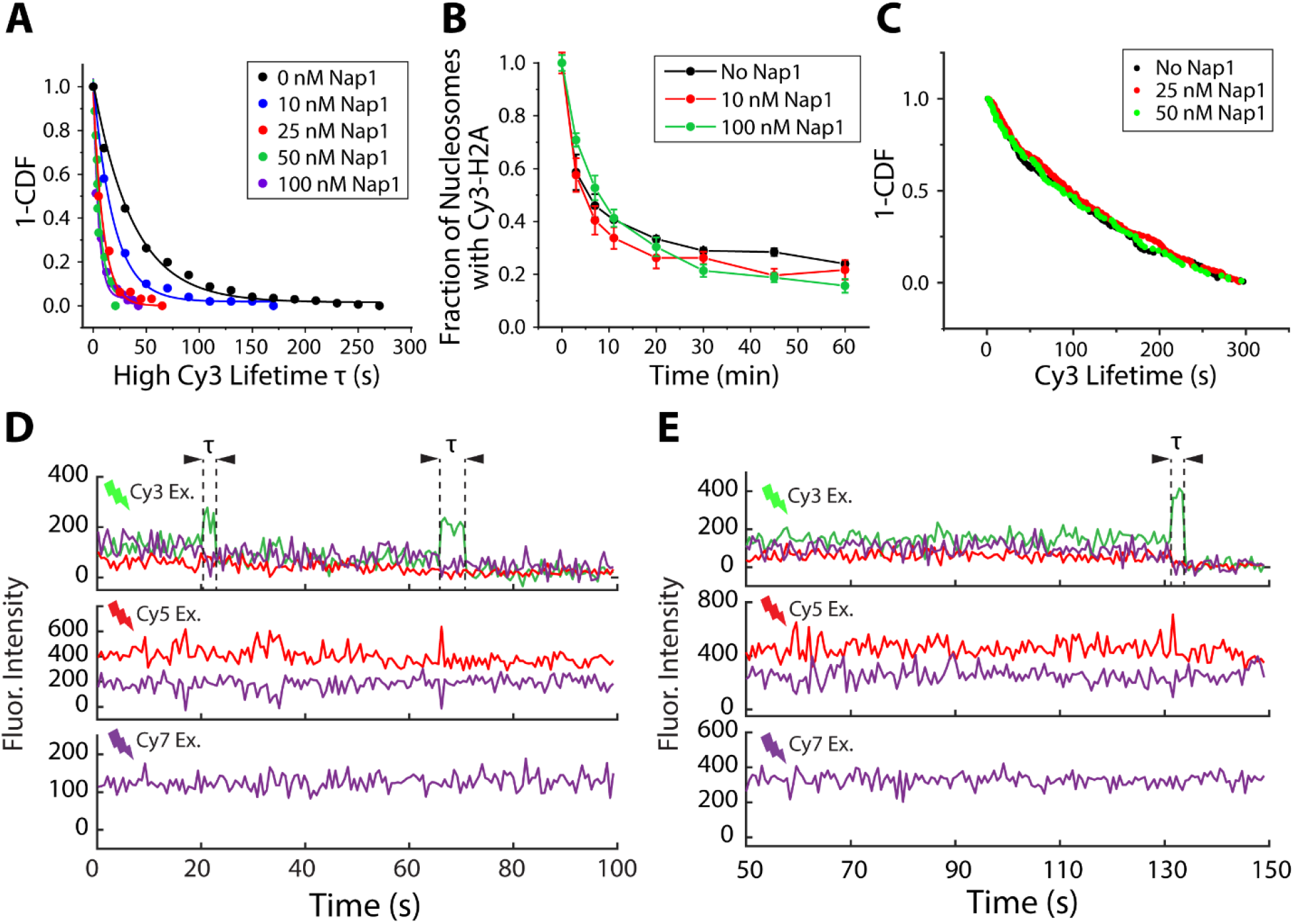
Nap1 removes H2A-H2B dimer from the SWR1-nucleosome complex. (**A**) 1-CDF of the high Cy3 lifetime (τ) under different Nap1 concentrations. Lines are single exponential fits to the data. Results were pooled from two technical replicates. (**B**) The Cy3-AB eviction rate (as measured in Fig. 1D) in the presence of various Nap1 concentrations. Nap1 does not significantly change the rate of AB eviction. Results were averages of two technical replicates. (**C**) The Cy3 signal lifetime for all nucleosomes (exchanged or unexchanged) when 0, 25 nM or 50 nM Nap1 is present in the histone exchange reaction. Fitting 1-CDF to single exponential decay (R^2^ = 0.99) reveals the Cy3 lifetime is 145 ± 15 s when no Nap1 is present, 180 ± 18 s when 25 nM Nap1 is present and 152 ± 21 s when 50 nM Nap1 is present. These lifetimes were similar, suggesting Nap1 does not affect the photostability of Cy3. (**D, E**) Single-molecule time traces showing histone eviction when (**D**) 10 nM or (**E**) 50 nM Nap1 is present in the reaction chamber. The high Cy3 lifetimes here are noticeably shorter than when Nap1 is absent as shown in Fig. 2 and fig. S3.

**Fig. S6.**
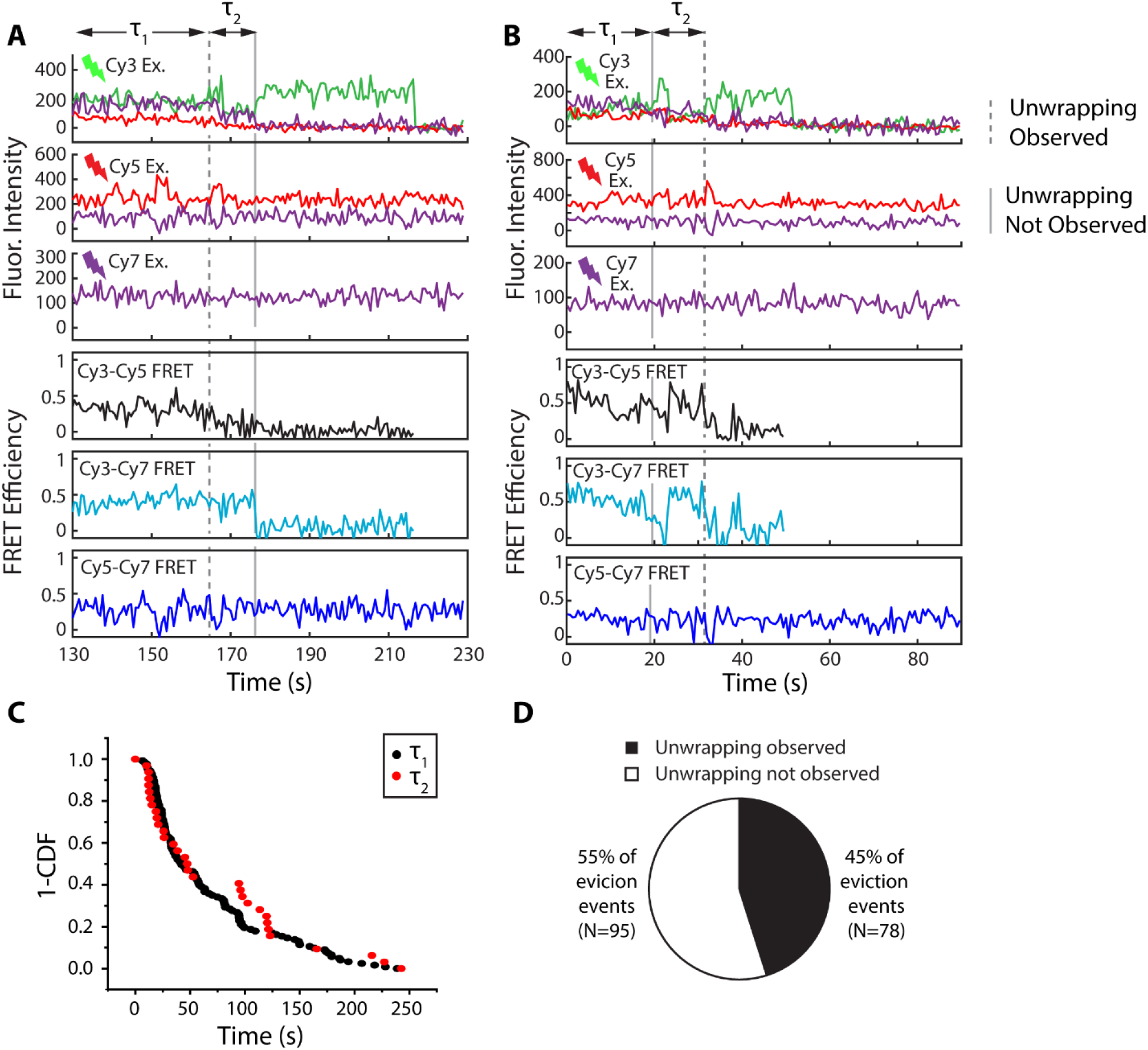
Sequential histone eviction can be observed in real-time. (**A,B**) Representative single-molecule time traces where two histone eviction events are captured. (**A**) A representative trace with two histone eviction events where the first histone eviction event (grey dashed line) shows a transient decrease in Cy5-Cy7 FRET (dark blue trace) while the second eviction event is not coincident with a Cy5-Cy7 FRET change (grey solid line). The second histone eviction event shows low Cy3-Cy5 FRET efficiency (black trace) prior to eviction, consistent with AB being evicted from the nucleosome face distal to Cy5. Therefore, the first histone eviction event must correspond to AB eviction from the proximal nucleosome face, the face where our assay is sensitive to DNA unwrapping. Indeed, we observed a transient decrease in Cy5-Cy7 FRET (blue) during the first eviction event. (**B**) Shows the opposite scenario, where the first eviction event (grey solid line) does not show a transient drop in Cy5-Cy7 FRET (dark blue), but the second eviction event does (grey dashed line). In this case, the second eviction event shows higher Cy3-Cy5 time-averaged FRET (black) prior to eviction, consistent with the Cy5-proximal AB being evicted. It follows that the first eviction event corresponds to Cy5-distal AB being evicted from the face where our assay is not sensitive to DNA unwrapping. Indeed, the Cy5-Cy7 FRET (dark blue) efficiency did not change during the first eviction event. t_1_ is defined as the time between injection of the reaction mixture at 0 s and the increase in Cy3 signal due to histone eviction. t_2_ is the time between the first and the second histone eviction events. (**C**) 1-CDF (cumulative distribution function) of t_1_ and t_2_. Each data point corresponds to a single measurement. t_1_ = 58.1 ± 4.5 (R^2^ = 0.99), t_2_ = 57.5 ± 14.6 s (R^2^ = 0.96). t_1_ and t_2_ were the same, suggesting that SWR1 is not processive. (**D**) The fraction of histone eviction events coincident with DNA unwrapping.

**Fig. S7.**
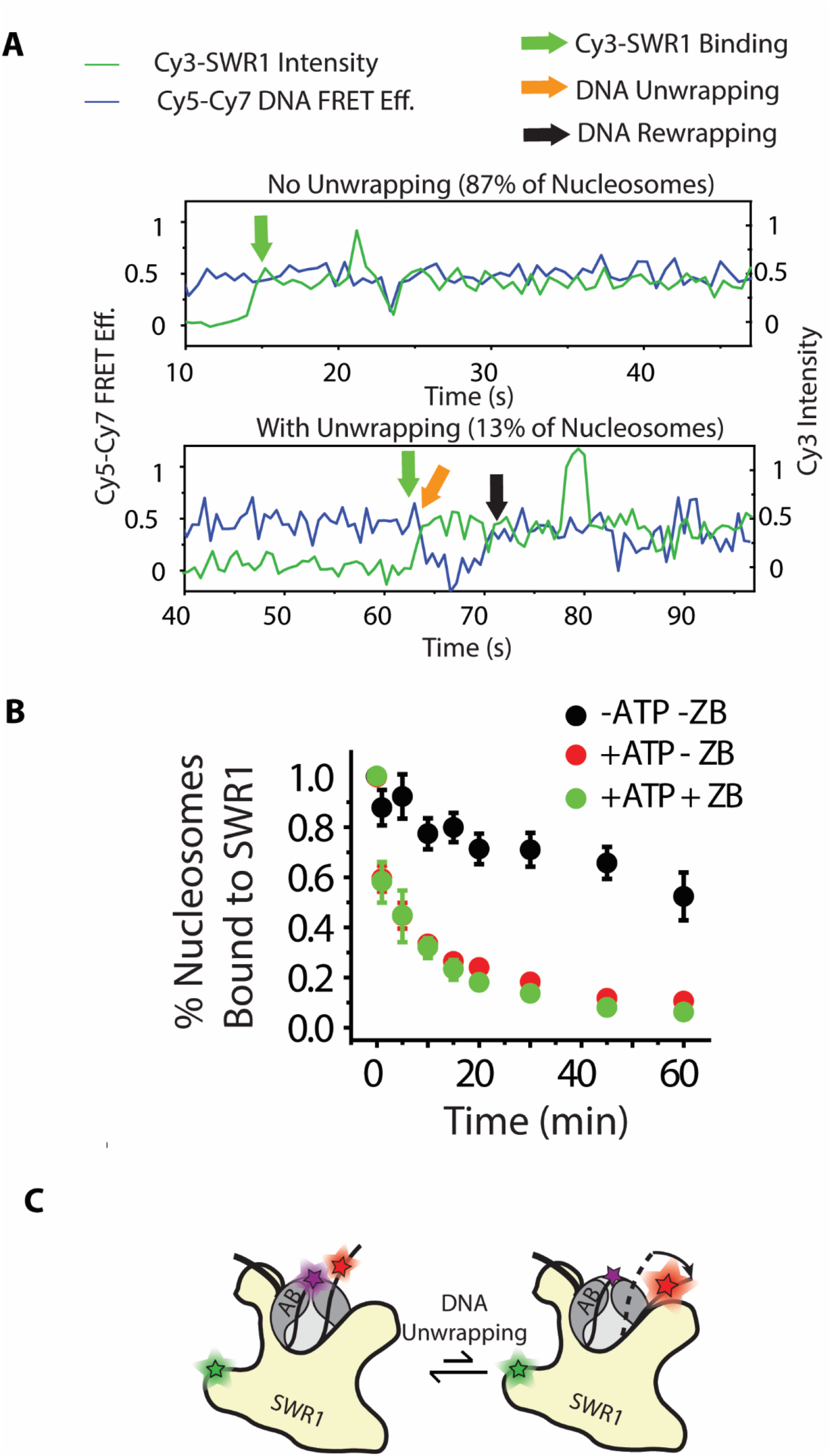
Nucleosomal DNA remains wrapped when SWR1 is bound. (**A**) Cy3 intensity (green line) and Cy5-Cy7 FRET (blue line) trajectories for two nucleosomes labeled with Cy5 and Cy7 on the nucleosomal DNA and exposed to 5 nM Cy3-labeled SWR1. The top trace is representative of a nucleosome that did not unwrap upon SWR1 binding (green arrow). This is representative of the majority (87%) of traces showing that nucleosomal DNA remains wrapped even while SWR1 is bound. The bottom trace is representative of a nucleosome that did unwrap (orange arrow) and rewrap (black arrow) during the time SWR1 was bound. This trace is representative of only 13% of SWR1-bound nucleosomes. (**B**) Measuring the fraction of nucleosomes that are Cy3-SWR1-bound after introducing 5 nM Cy3-SWR1 into the imaging chamber followed by a buffer wash to remove unbound Cy3-SWR1. This data shows that the SWR1-nucleosome complex is long-lived and, in these conditions, has a lifetime of tens of minutes if washed out with buffer alone. Interestingly, the addition of either 1mM ATP or 1 mM ATP and 10 nM ZB dimer into the wash buffer decreases the lifetime of the bound state dramatically. Data is mean of three different experiments +/− standard deviation. (**C**) A cartoon depicting that nucleosomal DNA remains wrapped most of the time SWR1 is bound, transiently unwrapping only for a short period of time.

**Fig. S8.**
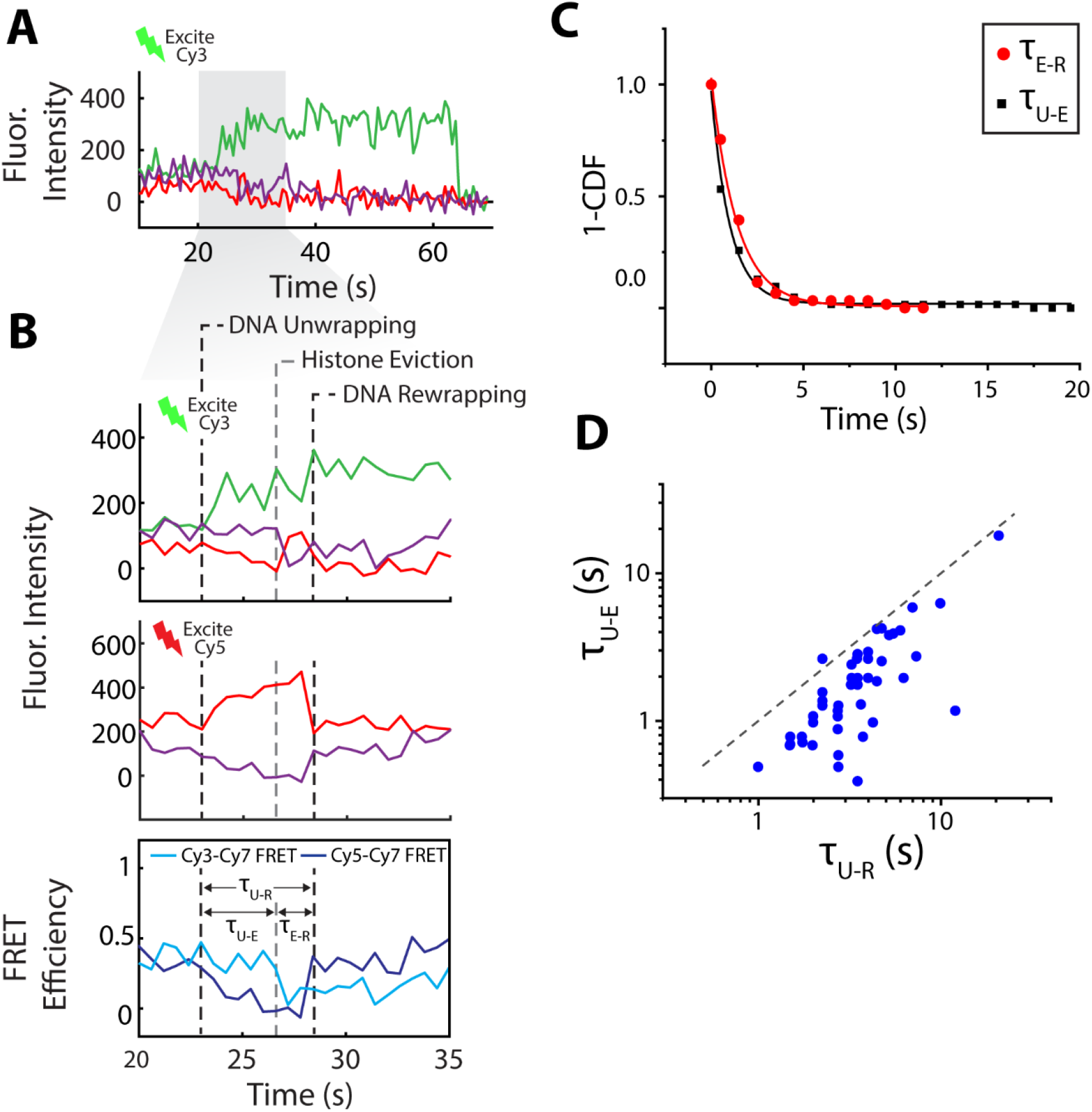
Additional single-molecule time traces and statistics on reaction intermediates during productive DNA unwrapping. (**A**) A representative single-molecule time trace showing histone eviction. (**B**) Expanded view of the shaded region in A, showing DNA unwrapping, histone eviction and DNA rewrapping can be distinguished via FRET. These events occur sequentially during the histone exchange reaction. (**C**) 1-CDF of τ_E-R_ and τ_U-E_ are fit to single exponential decays. τ_U-E_ = 1.01 ± 0.06 s, R^2^ = 0.99. τ_E-R_ = 1.39 ± 0.10 s, R^2^ = 0.99. (**D**) Scatter plot of paired τ_U-R_ and τ_U-E_ where each data point corresponds to measurements from a single histone eviction event. This plot shows that the time for histone eviction after productive DNA unwrapping (τ_U-E_) is always shorter than the time between the start of productive unwrapping and rewrapping (τ_U-R_). Therefore, H2A-H2B is always displaced from the histone octamer after DNA is first productively unwrapped, but before DNA is rewrapped, during the exchange reaction.

**Table S1.**
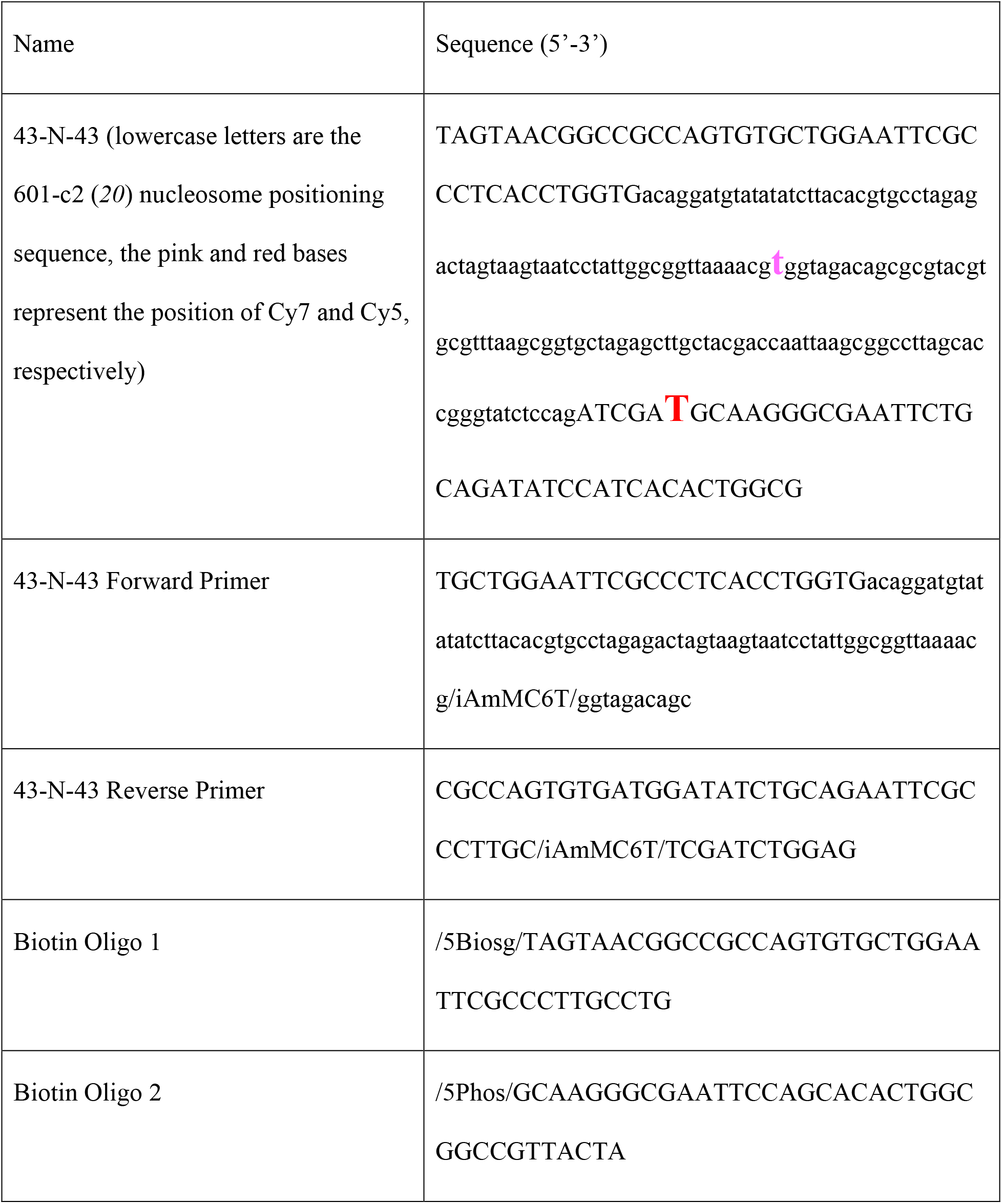
DNA and Primer Sequences for the biotin-43-N-43 Construct.

**Table S2.**
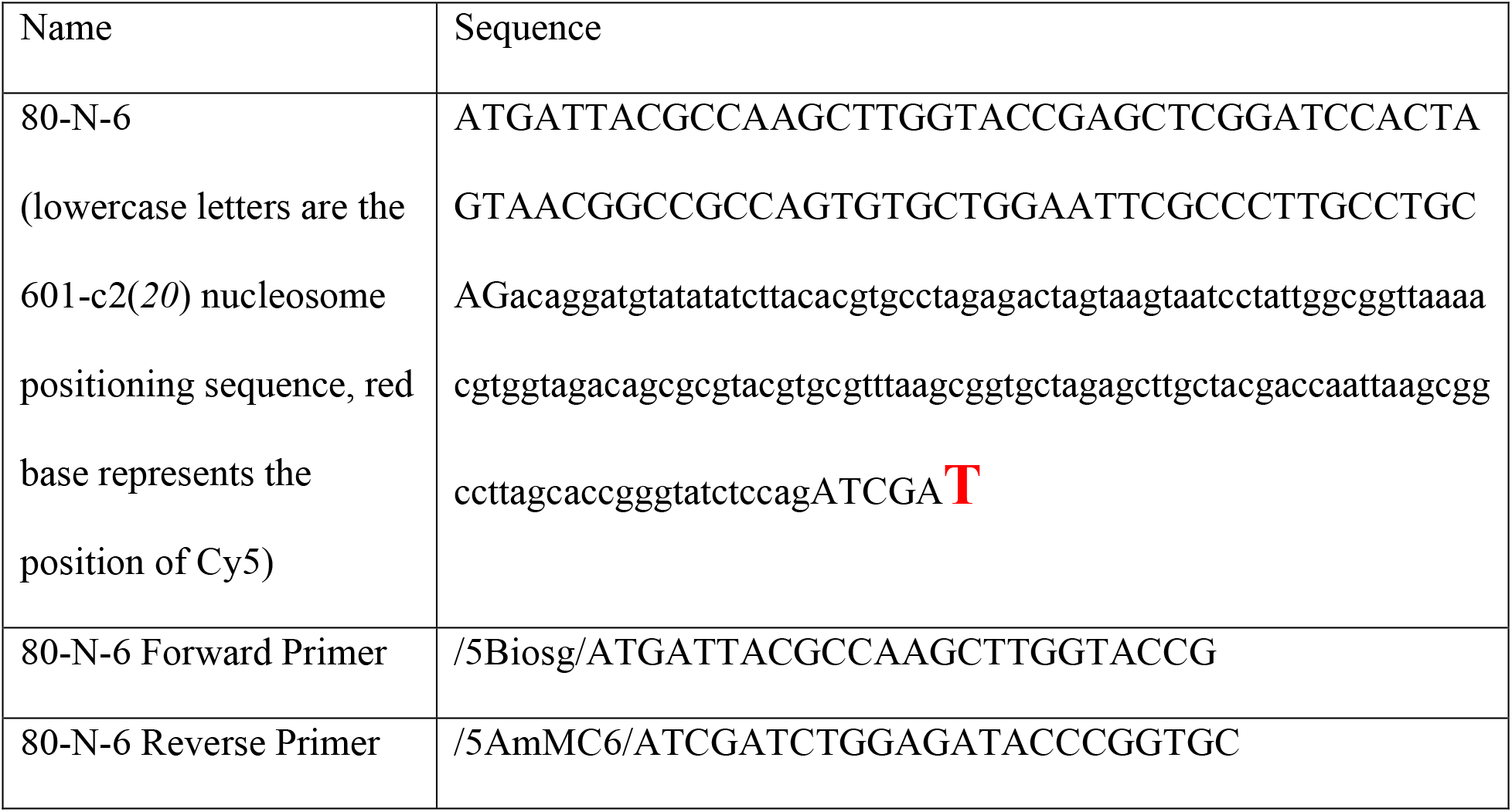
DNA and Primer Sequences for the biotin-80-N-6 Construct.

**Table S3.**
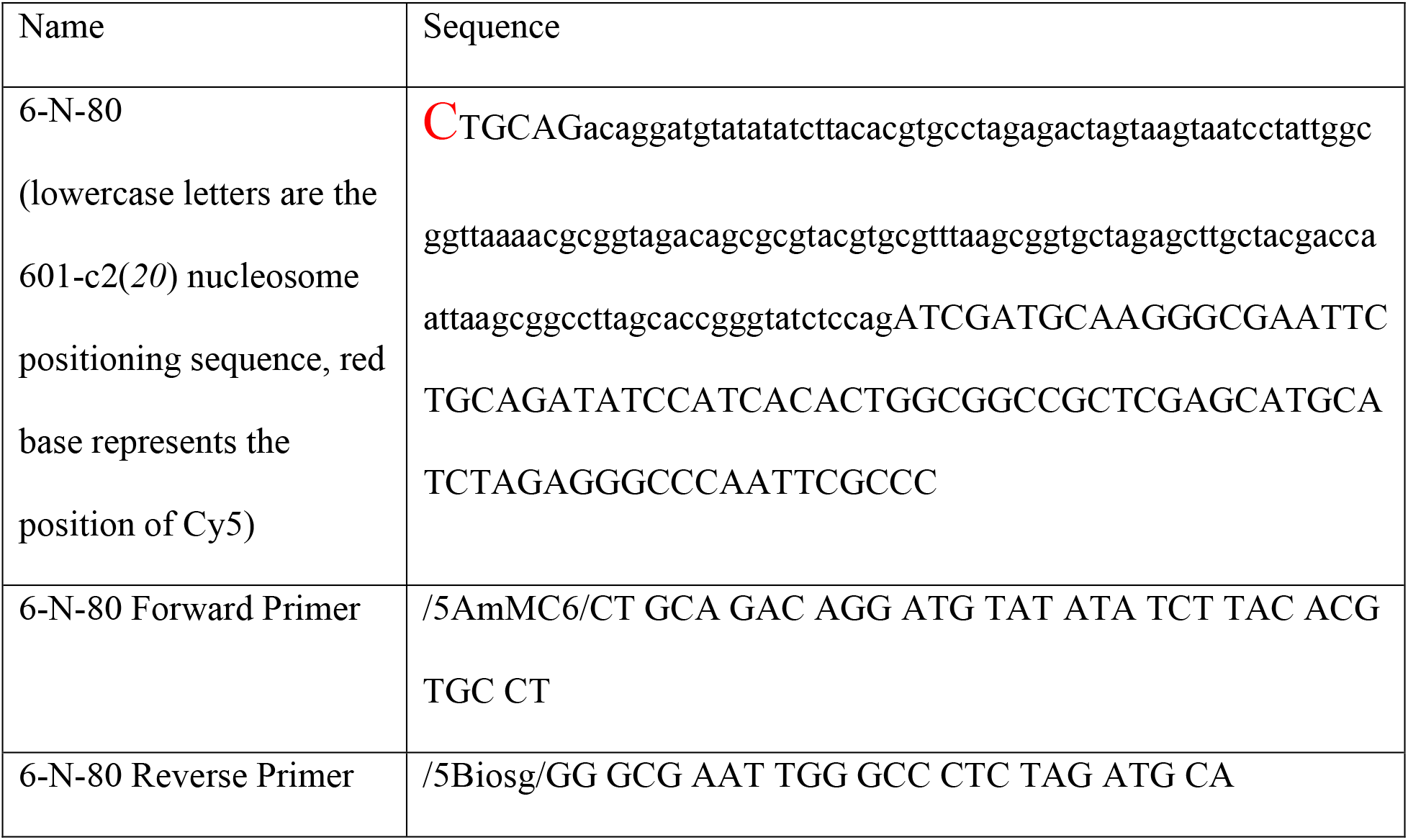
DNA and Primer Sequences for the 6-N-80-biotin Construct.

